# Stochastic activation and bistability in a Rab GTPase regulatory network

**DOI:** 10.1101/776567

**Authors:** Urban Bezeljak, Hrushikesh Loya, Beata Kaczmarek, Timothy E. Saunders, Martin Loose

**Author notes:** Authors for correspondence: M.L.; T.E.S.

## Abstract

Rab GTPases are the central regulators of intracellular traffic. Their function relies on a conformational change triggered by nucleotide exchange and hydrolysis. While this switch is well understood for an individual protein, how Rab GTPases collectively transition between states to generate a biochemical signal in space and time is unclear. Here, we combine in vitro reconstitution experiments with theoretical modeling to study a minimal Rab5 activation network. We find that positive feedback in this network gives rise to bistable switching of Rab5 activation and provide evidence that controlling the inactive population of Rab5 on the membrane can shape the network response. Together, our findings reveal new insights into the non-equilibrium properties and general principles of biochemical signaling networks underlying the spatiotemporal organization of the cell.

## Introduction

Positive feedback is a core motif in biochemical circuits that can generate bistable behavior, where the system can collectively switch between an ON and OFF state (1). Regulatory networks incorporating positive feedback loops control various cellular processes, such as cell polarization (2), oocyte maturation (3), and cell cycle progression (4). Positive feedback has also been proposed to be important for the organization of membrane traffic by small GTPases (5–7). Despite such ubiquity, the molecular events underlying the emergent properties of these networks are currently poorly understood.

Small GTPases of the Rab family organize the eukaryotic endomembrane system by defining the biochemical identities of organelles and directing membrane traffic between intracellular compartments through vesicle formation, transport, docking, and fusion with the target organelle (8). Arguably the best characterized Rab GTPase is Rab5, which controls the maturation of early endosomes towards the lysosomal system (9). Like all small GTPases, Rabs can exist in either an active GTP- or inactive GDP-bound state. Additionally, Rab GTPases possess one or two lipophilic geranylgeranyl chains on their C-terminal, which anchor them to the membrane surface (10). There, they recruit downstream effectors to orchestrate the vesicular flow. The transition between nucleotide states is controlled by guanine nucleotide exchange factors (GEFs) that catalyze exchange of GDP with GTP; and GTPase activating proteins (GAPs) catalyzing GTP hydrolysis (11). In their inactive GDP-bound state, the Rab GDP-dissociation inhibitor (GDI) extracts the Rab GTPase from the membrane and keeps it soluble in the cytoplasm (12). As a result, nucleotide exchange and hydrolysis drive dynamic cycling of the GTPase to and from the membrane. In the case of Rab5, the GEF Rabex5 forms a complex with the Rab5 effector Rabaptin5 (13). Consequently, Rab5 is thought to recruit its own activator to establish a positive feedback motif, which was proposed to result in its ultrasensitive activation (14) and membrane accumulation (13, 15–19). However, whether these molecular interactions can indeed lead to switch-like activation and collective membrane binding of Rab5 is not known (20).

The reason for this lack of understanding is that the characterization of small GTPase networks on a systems level has remained challenging. First, the inherent complexity of the living cell makes *in vivo* control over reaction conditions and precise experimental readouts challenging. Second, in contrast to the situation *in vivo*, activity studies performed *in vitro* commonly relied on proteins without their physiological geranylgeranyl modifications and were performed in the absence of the GDI and membranes (17, 21). Accordingly, these simplified experimental setups can lead to non-physiological activation dynamics (22). Lastly, the input-output relationship of the Rab GTPase activation switch in a biologically relevant setting is currently unknown as previous *in vitro* assays of Rab regulation did not address the non-equilibrium dynamics of small GTPases under cycling conditions (15, 23, 24).

Here, we rebuild the dynamic network underlying Rab5 activation *in vitro* using a minimal set of purified components (Fig. 1 and S1): fluorescently labeled, prenylated Rab5 in complex with GDI; Rabex5:Rabaptin5; and biomimetic membranes. In combination with theoretical modeling, this experimental approach allowed us to assay Rab5 activation far from biochemical equilibrium and to study the mechanisms of collective Rab5 activation under controlled conditions.

**Fig. 1.**
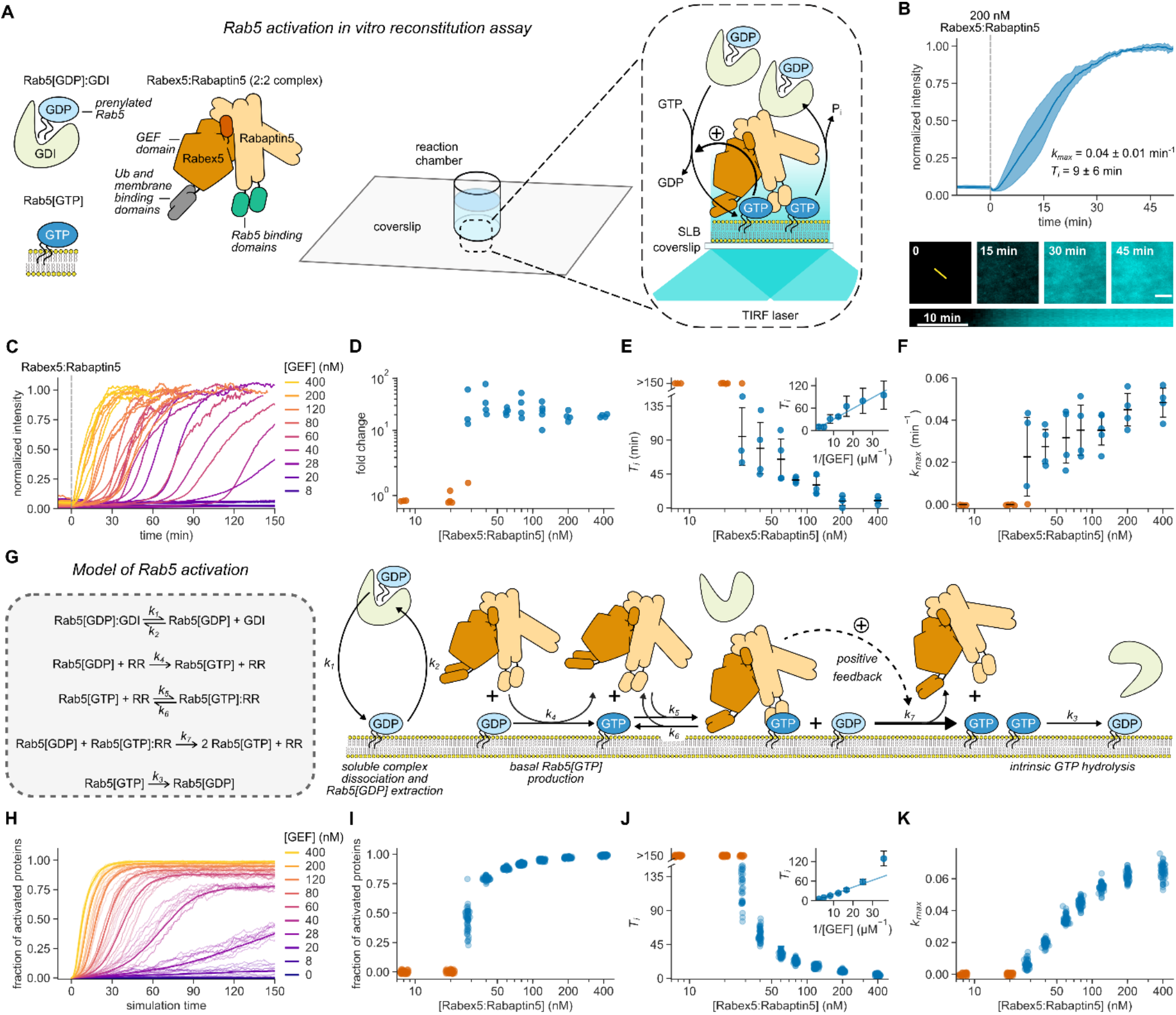
Rab5:GDI activation on phospholipid membranes is ultrasensitive and stochastic. **(A)** Schematic of Rab5 activation reconstitution assay on a supported lipid bilayer (SLB). **(B)** Top panel: addition of Rabex5:Rabaptin5 triggers nucleotide exchange by CF488A-Rab5, which can be followed by an increase of fluorescence intensity on the membrane surface. Solid line is mean normalized intensity, shaded area corresponds to SD (n = 4). Bottom: micrographs of CF488A-Rab5 binding to the SLB after addition of 200 nM GEF complex and corresponding kymograph (below) taken along the yellow line. Scale bar = 5 µm. **(C)** Rab5 intensity traces obtained at increasing Rabex5:Rabaptin5 concentrations. **(D)** Rab5:GDI-Rabex5:Rabaptin5 activation response curve. The fold change was calculated by dividing the fluorescence intensity at steady state with the average signal 10 min before GEF addition. **(E)** Activation delay *T*_*i*_ decreases with higher Rabex5:Rabaptin5 concentration. Where no detectable activation was observed within 150 min, the *T*_*i*_s are denoted as >150 min and shown in orange. Error bars are SD. **(F)** Relative maximum rates *k*_*max*_ against the GEF complex concentration reveal cooperativity of Rab5 activation. Without detectable activation within 150 min, the activation rate was determined to be 0 and the corresponding points are depicted in orange. Error bars are means ± SD. **(G)** Schematic representation of modeled molecular interactions. We constructed a model of the minimal Rab5 activation network based on the known literature (13, 15–19, 38). We then derived ODEs based on mass action kinetics. **(H)** Stochastic model simulations of Rab5 activation at increasing Rabex5:Rabaptin5 particle numbers. Shown are average curves from 50 individual runs in bold and 10 random traces per condition. **(I** – **K)** Signal fold change, temporal delays and relative maximum rates from the stochastic simulations in (H). We ran 50 individual stochastic simulations per condition.

## Results

First, we set out to verify the activity of purified Rabex5:Rabaptin5 on Rab5[GDP] in complex with GDI. We loaded lipid-modified Rab5 with the fluorescent GDP analog mant-GDP and used its fluorescence intensity as a real-time readout of nucleotide exchange (13, 16, 17). With 60 nM GEF and in the absence of membranes, we could not detect nucleotide exchange on 250 nM Rab5[mant-GDP]:GDI. However, we found robust activation in the presence of small unilamellar vesicles (SUVs) (Fig. S2), confirming that the phospholipid bilayer is essential for activation of the Rab:GDI complex (25, 26). To investigate the role of biological membranes for Rab5 activation, we utilized glass supported lipid bilayers (SLBs) as membrane substrates, combined with fluorescently labeled proteins and TIRF microscopy (Fig. 1A) (27). To recapitulate the intracellular pre-activation state, we first incubated the SLB with inactive CF488A-Rab5:GDI (500 nM), 0.5 mM GTP and 0.05 mM GDP. We included free GDI (2 µM) to mimic cellular stoichiometric excess of RabGDI (28). We then initiated nucleotide exchange by adding 200 nM Rabex5:Rabaptin5 and followed the fluorescence of CF488A-Rab5 on the membrane. Starting from low basal level of fluorescence on the membrane surface, the addition of the GEF complex produced a characteristic rise in fluorescence intensity until the signal saturated after about 40 minutes (Fig. 1B), consistent with an accumulation of Rab5[GTP] on the membrane. Accordingly, SLBs can act as a membrane substrate for prenylated Rab5, allowing us to follow its collective activation and membrane binding in real time.

Positive feedback regulation typically gives rise to sigmoidal signal-response curves (29). To test for the presence of positive feedback in the Rabex5:Rabaptin5:Rab5 activation network we recorded Rab5 membrane binding after adding increasing amounts of the GEF complex (Fig. 1C). Strikingly, we found that this titration resulted in an apparent two-state response profile: while there was no activation at GEF concentrations below 20 nM even 150 minutes after Rabex5:Rabaptin5 injection, we found a 10- to 80-fold increase of fluorescence on the membrane with higher concentrations of Rabex5:Rabaptin5 (Fig. 1D). From the temporal activation curves, we extracted the relative maximal rate of Rab5 activation (*k*_*max*_) as well as the time delay needed to reach this rate (*T*_*i*_) (30) (Fig. S3, Materials and Methods). High GEF complex concentrations (400 nM) gave rise to an immediate increase in Rab5 fluorescence intensity. At intermediate GEF concentrations, we observed nearly flat intensity profiles for up to 2 hours before collective Rab5 activation (Fig. 1E). At low GEF concentrations, we observed no response within the measurement window (orange circles, Fig. 1E). We also performed extended time recordings at 8 nM GEF and saw no response even after up to 12 hours (Fig. S4). Interestingly, the temporal delays needed to reach half activation increased linearly with the inverse of GEF complex concentrations (Fig. 1E, inset). Despite different delay times, all activation profiles had a similar sigmoidal shape (Fig. S5). By plotting *k*_*max*_ against GEF concentration, we found that nucleotide exchange showed high cooperativity (Fig. 1F) with a critical GEF concentration of around 28 nM, where we observed significant variations between the response curves, with some measurements having no significant response over the time course of the experiment. Below this point, no collective switching was detected, while higher GEF concentrations allowed for fast activation and Rab5 membrane accumulation, which gradually increased (17).

To better understand the dynamic response curves and the origin of the observed activation delays, we constructed a model of the minimal reaction network, which includes cooperative activation due to a direct interaction of Rab5[GTP] with its GEF complex (Fig. 1G, Supplementary Text). Precise details of this cooperative interaction are not known, so in the model we take a conservative approach whereby the positive feedback is relatively weak. Solving the model using the Gillespie algorithm to incorporate biochemical noise (stochasticity) in the reactions (31) produces similar dynamics and time delays to those observed experimentally (Fig. 1H-K). In the absence of stochasticity, the predicted response curves deviated from the experiments: (1) at early times the intensity profiles were not flat, unlike measured experimentally; and (2) near the critical Rab5 concentration (~30nM), the model cannot replicate the broad range of activation times (Fig. S6). We cannot discount potential variations (*e.g.* precise initial protein concentrations) between each experiment playing a role in the observed results. However, given the highly controlled nature of our reconstituted experiment, we expect these fluctuations to be small. Together, our experimental and theoretical results provide clear evidence for positive feedback within a minimal Rab activation network sufficient to generate switch-like, ultrasensitive behavior. Furthermore, stochasticity is relevant for the system response near the critical switching concentration.

What are the molecular interactions giving rise to the observed cooperativity? It has been proposed that cooperative Rab5 activation is due to GTP-dependent, effector-mediated GEF recruitment (13, 15–19). Alternatively, direct binding of Rabex5 to the negatively charged membrane could also enhance nucleotide exchange by retaining the GEF complex on the membrane (32). To test these possibilities, we prepared Δ_RBD_Rabaptin5, which lacks Rab5 binding domains (RBDs) (20); and ΔRabex5, which misses putative membrane targeting motifs (16) (Fig. 2A). Of all GEF complex variants tested, we detected efficient Rab5 activation only for full length Rabex5:Rabaptin5 and ΔRabex5:Rabaptin5. In contrast, there was no collective activation in the absence of Rabaptin5 (ΔRabex5) or for the GEF complex without the Rabaptin5 RBDs (Rabex5:Δ_RBD_Rabaptin5) (Fig. 2B). The same dependence on Rab5:Rabaptin5 interaction was also apparent in our model (Fig. 2C). Using fluorescently labeled Rabaptin5 and dual color imaging, we found that Rab5 and the GEF complex showed similar intensity traces in experiments (Fig. 2D) and in our model (Fig. 2E), confirming that Rabex5:Rabaptin5 is retained on the membrane surface by active Rab5[GTP] to engage the positive feedback loop (33, 34). Together, these results demonstrate that Rabaptin5 not only enhances the GEF activity of Rabex5 (17, 35), but that direct interactions between GTPase, GEF and effector in a ternary complex are essential for the cooperative activation of Rab5 and its collective binding to the membrane (20).

**Fig. 2.**
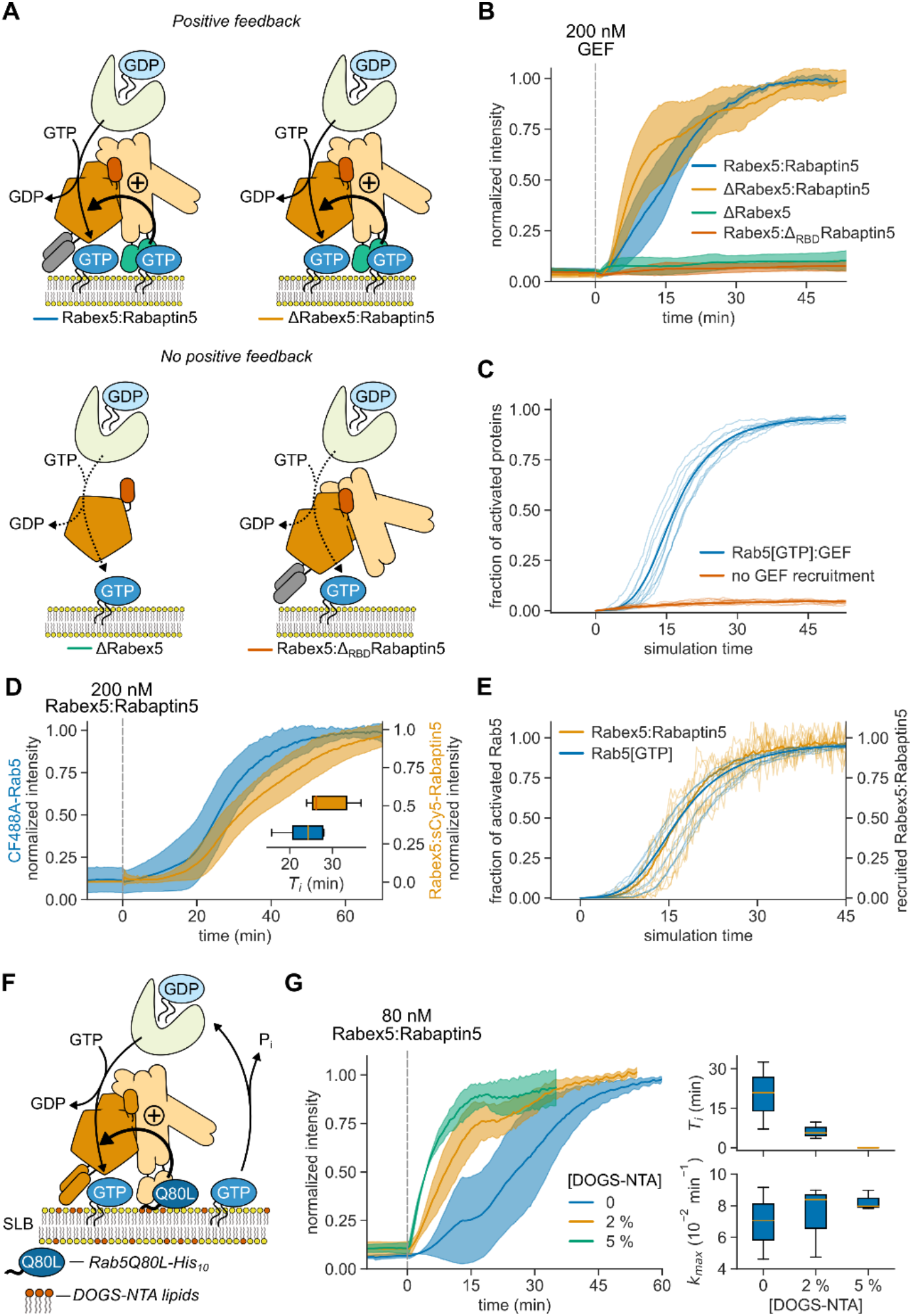
Positive feedback of Rab5 activation depends on GEF recruitment. **(A)** Illustration of protein interactions responsible for collective Rab5 switching. Positive feedback originates from a direct interaction between Rabex5:Rabaptin5 and Rab5[GTP]. **(B)** Fluorescence intensity traces obtained from experiments depicted in (A). Solid lines are mean normalized intensities, shaded areas are SD (Rabex5:Rabaptin5, ΔRabex5:Rabaptin5 n = 4; ΔRabex5, Rabex5:Δ_RBD_Rabaptin5 n = 3). **(C)** Stochastic model simulations with and without Rabex5:Rabaptin5:Rab5[GTP] complex formation (*k*_*5*_, *k*_*6*_ = 0) for 200 Rabex5:Rabaptin5 particles. Average curves from 50 individual runs are depicted in bold with 10 random traces per condition. **(D)** Kinetic traces of CF488A-Rab5 and Rabex5:sCy5-Rabaptin5 activation. Solid line is mean relative normalized fluorescence intensity, shaded area is SD (n = 5). Inset: *T*_*i*_ for CF488A-Rab5 (blue) and Rabex5:sCy5-Rabaptin5 (orange). **(E)** Stochastic model simulations for Rab5 and Rabex5:Rabaptin5 membrane binding for 200 Rabex5:Rabaptin5 particles. Shown are curves from 50 independent runs, the mean line is depicted bold with 10 random traces per condition. **(F)** Schematic of the reconstitution experiment with pre-activated SLB-immobilized Rab5Q80L-His10[GTP]. **(G)** Collective switching is faster with pre-activated Rab5. Left: Rab5 switching time courses in presence of 500 nM Rab5Q80L-His_10_ with increasing DOGS-NTA lipid concentration in the SLB. Solid line is mean normalized fluorescence intensity over time, shaded area is mean ± SD (n = 3). Right: corresponding time delays *Ti* and relative maximum rates *k*_*max*_.

What could explain the long delay times and stochastic switching observed at intermediate concentration of the GEF complex? Typically, long lag phases are related to processes that rely on random nucleation events that trigger phases of rapid growth (36, 37). Importantly, these lag phases can be dramatically shortened in the presence of seeds that trigger activation. To test this prediction, we attached different amounts of GTP-loaded constitutively active Rab5Q80L-His_10_ on SLBs with nickel-chelating lipids (DOGS-NTA) before adding 80 nM Rabex5:Rabaptin5 (Fig. 2F). Without pre-activated Rab5 on the membrane (0 [DOGS-NTA]), activation occurred 20 min after addition of this concentration of Rabex5:Rabaptin5. In contrast, the time delays with Rab5Q80L-His_10_ on 2 % [DOGS-NTA] membranes were 3-times shorter and completely absent with 5 % [DOGS-NTA] (Fig. 2G), while the maximal activation rates were not significantly changed. This data shows that membrane-bound Rab5[GTP] can act as a seed for Rab5 activation and membrane accumulation.

Next, we wanted to find out what could initiate the Rab5 activation switch in the absence of active protein on the membrane. As the presence of membranes is required to activate the Rab5:GDI complex, we predicted that inactive Rab5[GDP] existing on the membrane prior to addition of the GEF complex is the substrate for nucleotide exchange (22, 38). Indeed, with small amounts of sCy5-Rab5:GDI in a background of CF488A-labeled Rab5:GDI, we found individual sCy5-labeled proteins on the membrane even before adding Rabex5:Rabaptin5 (Fig. 3A). Using single molecule tracking, we found that non-activated sCy5-Rab5 diffused rapidly on the membrane and had a mean residence time of 0.3 ± 0.1 s (Fig. 3B). After addition of the GEF complex, we found a sudden increase in sCy5-Rab5 particle counts, along with a sigmoidal increase of membrane-bound CF488A-Rab5. The histogram of membrane residence times of Rab5[GTP] and corresponding fits revealed two populations: a short-lived population with a residence of 0.4 ± 0.2 s, similar to Rab5[GDP], and a long-lived population with a 10-times longer residence time (3.3 ± 1.3 s) (Fig. 3B). A similar membrane lifetime distribution was observed for Rab5 with the non-hydrolyzable GTP analog GMP-PNP (Fig. S7) indicating that the values for activated Rab5 are influenced by fluorophore bleaching and represent a lower bound of membrane-residence time. Together, these results indicate that Rab5 first transiently binds to the membrane in its GDP-bound state, before it is converted by Rabex5:Rabaptin5 to its long-lived GTP-bound state. Rab5[GTP] on the membrane can then act as seed that retains GEF complex and initiates the positive feedback. Accordingly, initial random activation events are likely the cause of the observed stochasticity for its collective transition to the active state.

**Fig. 3.**
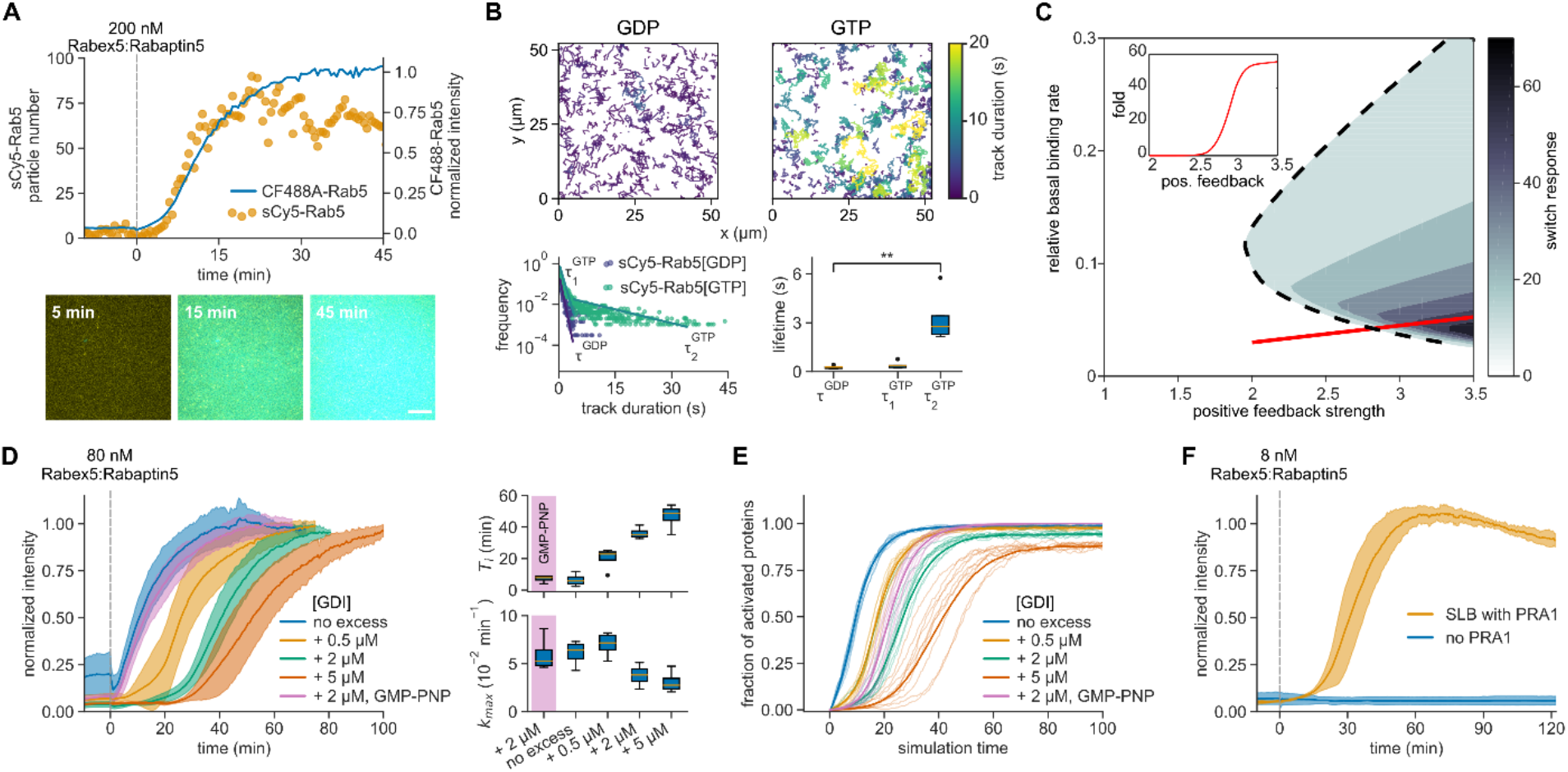
Rab5:GDI activation is tuned by free Rab5[GDP] abundance. **(A)** Rab5 cycles between the membrane and solution before and after nucleotide exchange. Top: sCy5-Rab5 molecule counts per frame and collective CF488A-Rab5 activation. Bottom: snapshots of the activation reaction. sCy5- and CF488A-Rab5 are depicted in yellow and cyan, respectively. Scale bar is 10 µm. **(B)** Rab5 single molecule trajectories reveal GDP- and GTP-bound proteins on the membrane. Top: 500 tracks of membrane-bound sCy5-Rab5 particles before (GDP) and after (GTP) activation. Bottom: frequency histogram identifies two populations with distinct lifetimes. A monoexponential decay before activation with lifetime τ^GDP^ and two-exponential decay with lifetimes τ_1_^GTP^ and τ_2_^GTP^, respectively (n = 5). **(C)** Parameter phase space of the phenomenological model for Rab5 switching, depending on the basal rate of activation (a_0_/a_2_K) and the strength of positive feedback (a_1_/a_2_K). Switching is defined as the relative difference in steady-state concentration relative to the scenario with no positive feedback. Inset: fold activation along the red line in the diagram. Stochasticity introduced by solving the phenomenological model within a Fokker-Planck framework. See text for parameter definitions. **(D)** Stoichiometric GDI excess over Rab5 affects delay of Rab5 activation *in vitro*. Left: solid lines are mean normalized intensities over time, shaded areas correspond to SD (n = 3). Right: corresponding activation *T*_*i*_ and relative maximum rates *k*_*max*_. **(E)** Stochastic simulations of the full model for varying initial amounts of GDI excess (0 – 2000 particle number). Shown are curves from 10 random runs per condition, the mean line from 50 runs is depicted bold. **(F)** SLB-bound PRA1 enhances Rab5 activation at low GEF concentrations. Solid lines are mean normalized fluorescence intensities, shaded areas correspond to SD (n = 3).

How do the initial levels of membrane-bound Rab5 and the strength of the positive feedback affect the transition between the ON and OFF states? To answer this question, we used a coarse-grained (phenomenological) version of our model, which incorporated only binding (*a*_*0*_) and unbinding (*a*_*2*_) of Rab5 [R] on the membrane along with positive feedback (*a*_*1*_, with activation concentration *K*) (Fig. S8, Supplementary Text): 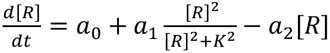. The parameter space that leads to GTPase switching (Fig. 3C and S9) reveals that the switch response (*i.e.* the fold change in membrane-bound Rab5) after activation is small when the basal biding rate is set high. Conversely, if the basal binding rates are too low, the critical threshold for switching fails to occur, even with stochastic fluctuations. This reveals that the system switching is potentially highly tunable, and dependent on both the basal binding rate and positive feedback strength.

To experimentally test the model predictions for how GTPase activation is tuned, we first varied the rate of extraction of Rab5[GDP] by adding different amounts of free GDI in our experiments (Fig. 3D). We found that increasing the stoichiometric GDI excess lowered the basal background fluorescence prior to activation, prolonged activation delay times after GEF addition, and limited *k*_*max*_, consistent with a decreased basal binding rate. Using our full model, we also see similar results when altering the level of free GDI (Fig. 3E) confirming that high membrane extraction rates of Rab5[GDP] cause long delay times and stochastic activation (39).

To increase basal Rab5 binding, we first replaced GTP in our experiment with GMP-PNP. As this GTP analog inhibits Rab5’s high intrinsic GTPase activity (40), it should prevent extraction of activated Rab5 from the membrane and therefore lead to a more robust transition into the ON state. In agreement with this prediction, we observed immediate collective Rab5 membrane binding after adding 80 nM GEF complex with GMP-PNP and 2 µM GDI, while the delay time was more than 36 min when we used GTP (Fig. 3D, magenta curve). Preventing Rab5 membrane extraction in the full model but keeping other parameters fixed, we see that our model displays similar behavior for the GMP-PNP nucleotide exchange (Fig. 3E, magenta curve). Next, we added the Rab5-specific GDI dissociation factor – PRA1 to our experiments, which has been suggested to accelerate the release of Rab[GDP] from the GDI complex (41). Accordingly, it should also increase the basal GTPase binding rate and facilitate the collective activation switch. Indeed, with PRA1 in the membrane, we observed fast Rab5 activation with short delay times even at a Rabex5:Rabaptin5 concentration too low to support Rab5 activation on PRA1-free membranes (8 nM) (Fig. 3F). These findings show that despite not strictly required for Rab5 activation (38, 42), the presence of PRA1 in the endosomal membrane can lower the threshold for positive feedback initiation, making collective Rab5 activation more likely.

Conversely, further increasing Rab5’s GTPase activity above its intrinsic rate should inhibit collective switching as it prevents effector recruitment of the GEF complex and facilitates extraction of Rab5 from the membrane (Fig. 3C; moving to the left along the red line). To test this prediction, we performed experiments in the presence of purified full-length RabGAP-5 (SGSM3), a Rab5-specific GAP (43), which stimulates GTP hydrolysis by Rab5. We recorded the signaling response after addition of 80 nM Rabex5:Rabaptin5 in the presence of increasing RabGAP-5 amounts (Fig. 4A) and found that while it increased the activation delay, it did not substantially affect the maximal rate of Rab5 activation (Fig. 4B and 4C). At RabGAP-5 concentrations between 100 and 250 nM the reconstituted network either showed successful activation events or no accumulation of Rab5 on the membrane for different replicates at identical initial conditions. Importantly, once the system was switched ON, we found that even addition of 2 μM GAP (Fig. 4D) does not completely reverse the system to its pre-activated state. Similarly, by increasing the dissociation rate *a*_*2*_ in our phenomenological model, we observed clear difference in switching responses after 150 minutes, depending on the initial state of the network (Fig. 4E). This hysteretic response of the system confirms the bistable behavior of the Rab5 activation network.

**Fig. 4.**
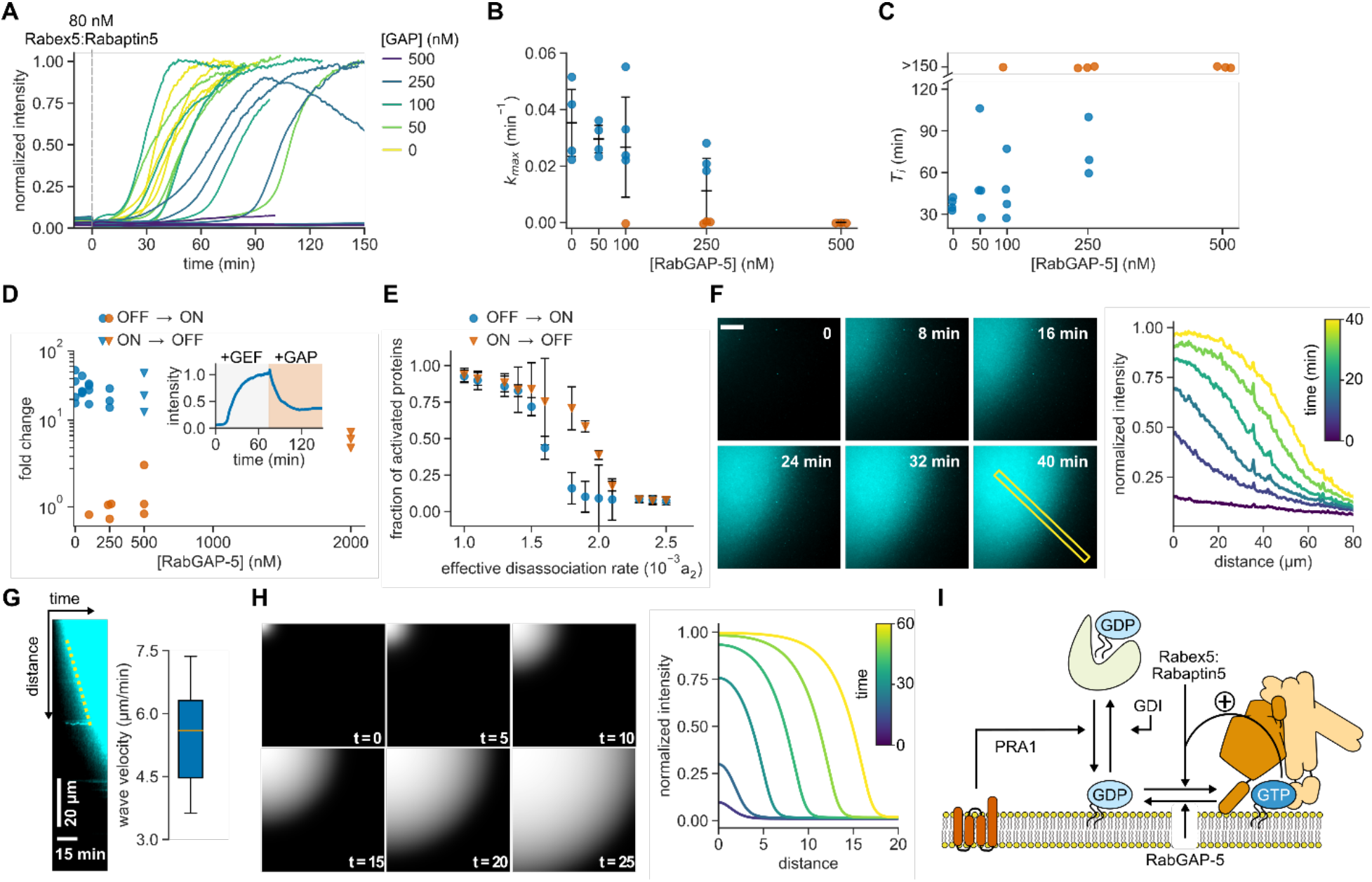
GAP activity reveals bistability of the reconstituted network. **(A)** Effect of RabGAP-5 on Rab5 activation. Shown are time courses at increasing GAP concentrations. **(B)** Maximal rates *k*_*max*_ of Rab5 activation for curves shown in (A). Without detectable activation within 150 min, the activation rate was set to 0 and the corresponding points are depicted in orange. Error bars are SD. **(C)** Activation delay *T*_*i*_ for data presented in (A). Without detectable activation, the times to inflection point are denoted as >150 min (orange). **(D)** GAP titration response curve. The fold change was calculated by dividing the fluorescence intensity at steady state with the average fluorescence signal 10 min before GEF addition. For ON → OFF switching, the system first reached active state (ON) with 80 nM GEF. Then, RabGAP-5 was added and the reaction was followed until the system reached a new steady state (OFF). Inset: ON → OFF switching time course with 2 μM RabGAP-5. **(E)** Changing the dissociation rate reveals hysteresis in switching of the phenomenological model after 150 minutes. Shown are means of 20 simulations, error bars are ± SD. **(F)** Left: Rab5 activation wave spreading across the SLB. Scale bar is 20 µm. Times indicate relative duration after start of acquisition, not time after addition of GEF complex. Right: fluorescence intensity profile of the indicated area. **(G)** Kymograph of the indicated area in (F) and mean wave velocity. Wave velocity was determined from the slope of fluorescence increase in generated kymographs (n = 6). **(H)** Simulated Rab5 activation front from including diffusion, *D*, within the phenomenological model; see Supplementary Text Eq. 6. Solution in terms of the dimensionless distance 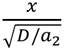. **(I)** Overview of the reconstituted Rab5 network regulation.

Strikingly, at RabGAP-5 concentrations of 50 nM, we observed Rab5 activation fronts on the membrane, where areas of high Rab5 density coexisted next to low Rab5 density areas (in 9 out of 13 experiments, in 3 experiments no obvious waves were noticed during activation, while in one experiment no activation occurred). This spatiotemporal activation pattern existed for more than 30 min, during which the activation front spread at a velocity of 5 μm/min before the system settled into a fully active state with Rab5 covering the SLB at high density (Fig. 4F, G). What could explain the emergence of this spatial pattern? Local activation of Rab5[GTP], due to random fluctuations, is reinforced and stabilized by positive feedback via engagement of Rabex5:Rabaptin5. This region of initial Rab5[GTP] activation will have higher probability of further Rab5[GTP] recruitment at its boundary than elsewhere on the surface, giving rise to a propagating activation front. This emergent property can be captured in our phenomenological model by introducing a diffusive term, where Rab5 activation in presence of a GAP can spread at constant rate by propagating the positive feedback activation via an activation front (44). Such a front is dependent on the GAP activity and the threshold Rab5[GTP] density that can sustain the positive feedback activation (Fig. 4H, Supplementary Text). It is well known that dynamic biochemical systems composed of locally acting cooperative actuators and long-ranged inactivators can give rise to chemical waves on the cellular and tissue level (45–48). In our system, RabGAP-5 acts as a global inhibitor, rather than a long-ranged diffusing inhibitor, resulting in our relatively simpler spatio-temporal patterns of activation.

## Discussion

To summarize, using *in vitro* reconstitution and theoretical modeling, we found that the minimal Rab5 regulatory network is ultrasensitive and bistable, likely prerequisites for the decisive signaling reactions controlling vesicle traffic. We have demonstrated that the architecture of the Rab5 activation network supports the formation of spatiotemporal patterns such as activation fronts, as found for other bistable systems with a local positive feedback and global inhibition (46). We also found that Rab5 activation in this minimal network can occur stochastically, and we identified the low amounts of non-active Rab5[GDP] as a potential source for this stochastic behavior. While stochasticity and long delay times are generally disadvantageous for intracellular signaling reactions that rely on tight control, our *in vitro* experiments demonstrate that it is possible to tune the response of the Rab5 activation network by regulating the stability of the Rab:GDI complex, either by the presence of a GDF in the membrane and possibly via GDI phosphoregulation (49).

Our study represents a systematic characterization of a minimal biochemical circuit of Rab GTPase activation. We have also provided examples for how additional regulatory interactions can be employed to direct and tune small GTPase activation in space and time. Of course, the composition of the cell provides more complex modes of regulation, both at the protein and membrane levels. Our *in vitro* system can be further extended to include other effectors or membrane compositions, making it an excellent testbed for probing the mechanisms of organelle identity formation during vesicle trafficking and the compartmentalization of the living cell. Furthermore, our approach can also be used to study the dynamic networks of other small GTPase families, such as Arf, Rac and Rho GTPases.

## Acknowledgments

The authors sincerely thank B. Simons, K. Kruse, M. Howard, E. Hannezo, A. Yap, T. Lecuit, and J. Brugués for discussions and valuable feedback on the manuscript. Additionally, we thank K. Loibl, other Loose lab members and the Scientific Service Units at the IST Austria for their support.

## Author contributions

Conceptualization, M.L.; Methodology, M.L., T.E.S. and U.B.; Software, T.E.S. and H.L.; Validation, U.B., H.L., T.E.S.; Formal Analysis, U.B., H.L. and T.E.S.; Investigation, U.B., B.K., H.L. and T.E.S.; Resources, U.B., B.K., H.L. and T.E.S.; Data Curation, U.B., H.L., T.E.S.; Writing – Original Draft, M.L., T.E.S. and U.B.; Writing – Review & Editing, U.B., H.L., B.K., T.E.S., M.L.; Visualization, U.B. and T.E.S.; Supervision, M.L. and T.E.S.; Project Administration, M.L. and T.E.S.; Funding Acquisition, T.E.S. and M.L.

## Data and materials availability

all data needed to evaluate and reproduce the reported conclusions is available in the manuscript or the supplementary materials.

