## Supplementary Information for "Stochastic activation and bistability in a Rab GTPase regulatory network"

**This PDF file includes:**

Materials and Methods
Supplementary Text
Figs. S1 to S9
Tables S1 to S3
Captions for Movies S1 to S6

**Other Supplementary Materials for this manuscript include the following:**

Movies S1 to S6

### Materials and Methods

Protein purification

**GDI.** *X. laevis* RabGDI was cloned as a $His_6$-(linker)-TEV-GDI construct in a pFastBac vector. The protein was expressed in Sf9 insect cell culture at 27 °C for 3 days after baculovirus infection. The harvested cell pellets were kept at -80 °C and thawed on the day of purification. The pellets were suspended in Lysis/Wash buffer (20 mM HEPES-NaOH pH 8.0, 500 mM NaCl, 5 mM 2-mercaptoethanol, 10 vol% glycerol) and supplemented with cOmplete EDTA-free protease inhibitor cocktail (Roche), 2.5 U/mL DNase I (Thermo Fisher Scientific) and 1 mM PMSF (Sigma Aldrich). We lysed insect cells using a glass douncer and 1 % IGEPAL CA-630 (Sigma Aldrich). The lysate supernatant was incubated with Protino Ni-IDA (Macherey Nagel) affinity resin after 45 min centrifugation at 60,000 rcf. After extensive washing of the affinity resin, we eluted the purified proteins with 150 mM imidazole in Lysis/Wash buffer. Next, we performed buffer exchange to the Lysis/Wash buffer using a PD-10 desalting column (GE Healthcare). The purification tag was later cleaved off with TEV(S219V) protease (pRK793; Addgene #8827) (*1*), which was removed by passing the solution over HisPur Ni-NTA resin (Thermo Fisher Scientific). Finally, the purified samples were concentrated on Vivaspin 20 concentrators (Sartorius), aliquoted and snap frozen in liquid nitrogen before storage at -80 °C.

**PRA1.** PRA1 was expressed for 4 h at 30 °C as TwinStrep-(linker)-TEV-$Gly_4$-PRA1 fusion in BL21(DE3) cells from a pET28a plasmid backbone. First, the cell pellets were resuspended in Lysis buffer (20 mM HEPES-KOH pH 7.8, 300 mM KOAc, 5 mM 2-mercaptoethanol, 10 vol% glycerol), supplemented with cOmplete EDTA-free protease inhibitor cocktail. The cells were lysed using cell disruptor (CF2, Constant Systems Ltd.) and 1 mM PMSF, 2.5 U/mL DNase I and 5 % Triton X-100 detergent (Sigma Aldrich) were added to the lysate. The transmembrane PRA1 was extracted with the detergent during 1 h incubation at 4 °C with mixing. We isolated the soluble fraction with 45 min centrifugation at 60,000 rcf and 4 °C. The supernatant was then loaded on StrepTrap HP affinity column (GE Healtcare), equilibrated with Purification buffer (20 mM HEPES-KOH pH 7.8, 300 mM KOAc, 5 mM 2-mercaptoethanol, 10 vol% glycerol, 0.1 % Triton X-100) and the bound proteins were eluted with 2.5 mM D-desthiobiotin (Sigma Aldrich). The pooled fractions were digested with TEV(S219V) protease overnight. The PRA1-detergent complexes were finally purified on Superdex 200 chromatography column (GE Healthcare). PRA1 suspension was stored at 4 °C for up to 5 days as freezing abolished the PRA1 activity. To produce PRA1-containing SLBs, 5 mM lipids with 79.9 % DOPC, 20 % DOPS and 0.1 % DMPE-PEG2000 (Avanti Polar Lipids) were dried under constant Nitrogen flow. Then, the lipids were hydrated in 100 μL Vesicle buffer (20 mM HEPES-KOH pH 7.4, 150 mM KOAc) with 20 mM Triton X-100. The purified PRA1 was added to the lipid suspension at 1 μM and the solution was incubated at room temperature for 10 min. We removed the detergent by chelation with 40 mM methyl-β-cyclodextrin (Sigma Aldrich) in Vesicle buffer, which was added to the suspension in 1:1 volume ratio. After 5 min incubation, the PRA1-lipid suspension was diluted to 0.5 mM and used to produce glass-supported membranes.

**Rab5.** A TwinStrep purification tag with TEV protease cleavage site and a $Gly_4$ linker was fused to codon-optimized *X. laevis* Rab5A. The whole construct was cloned into pFL baculovirus expression vector and expressed in Sf9 cells for 72 h at 27 °C. Notably, expression in alternative HighFive insect cell strain resulted in lower levels of post-translationally modified protein product. Rab5 was purified from insect cell pellets according to previously published protocol (*2*, *3*), with some modifications. First, the pellets were resuspended in Rab purification buffer (20 mM HEPES-KOH pH 7.4, 150 mM KOAc, 1 mM $MgCl_2$, 1 mM DTT and 10 vol% glycerol), supplemented with cOmplete EDTA-free protease inhibitor cocktail tablets, 2.5 U/mL DNase I and 1 mM PMSF. The cells were lysed by douncing and ten 1s pulses of tip sonication with 3s pauses. We isolated the prenylated Rab5 from the membrane fraction, which was collected by ultracentrifugation at 150,000 rcf for 30 min at 4 °C. We then resuspended the membrane fraction in Rab purification buffer, supplemented with 0.6 % CHAPS (Sigma Aldrich) and 100 µM GDP (Sigma Aldrich) (extraction buffer). The membrane-bound proteins were solubilized during 2 h incubation at 4 °C with agitation. The soluble fraction with extracted Rab5 was isolated by another ultracentrifugation at 150,000 rcf for 30 min at 4 °C. We loaded the supernatant on StrepTrap HP column in extraction buffer and bound proteins were eluted with 2.5 mM D-desthiobiotin. Next, the purification tag was cleaved with TEV(S219V) protease overnight. For fluorescent labeling, we used the sortagging method (*4*) with 10 µM Srt7M (Addgene #51141) (*5*) and 0.3 mM maleimide dye-CLPETGG peptide. For GEF activity assay, the unlabeled Rab5 was loaded with mant-GDP (Jena Bioscience) by 1 h incubation at 37 °C in presence of 2 mM EDTA and 20-times molar excess of the labeled nucleotide. The reaction was then quenched with 5 mM $MgCl_2$ and the buffer was exchanged back to extraction buffer using Zeba spin columns (Thermo Fisher Scientific). The Rab5:GDI complex was assembled by dialyzing the purified components in 1:2 molar ratio against the Rab purification buffer with 0.1 % CHAPS. If necessary, the dialyzed samples were concentrated on Amicon Ultra-2 filter units (Millipore). Finally, the complex was loaded on Superdex 75 size exclusion column (GE Healthcare) equilibrated with Rab purification buffer without any detergent. This separated the Rab5:GDI complex from individual components, the TEV protease, SrtA and unused labeled peptide. The peak fractions containing the Rab5:GDI complex were subsequently pooled and frozen in liquid nitrogen before storage at -80 °C.

**Rab5Q80L-$His_{10}$.** Unprenylated Rab5Q80L-$His_{10}$ from *X. laevis* was expressed in *E. coli* BL21(DE3) bacterial strain from pET28a vector. The protein was cloned as TwinStrep-(linker)-TEV-$Gly_4$-Rab5Q80L-$His_{10}$ fusion. The frozen bacterial pellets were thawed on ice and resuspended in Lysis/Wash buffer (50 mM HEPES-NaOH pH 8.0, 500 mM NaCl, 5 mM 2-mercaptoethanol, 10 vol% glycerol). To prevent non-specific protease digestion, cOmplete EDTA-free protease inhibitor cocktail tablets and 1 mM PMSF were added to the cell suspension. Next, the recombinant cells were lysed using cell disruption and centrifuged at 60,000 rcf for 45 min at 4 °C. The protein was purified using Protino Ni-IDA affinity resin and eluted in Lysis/Wash buffer with 150 mM imidazole. After buffer exchange on PD-10 columns to MonoQ A buffer (50 mM Tris-HCl pH 8.8, 1 mM DTT), the protein was digested with TEV(S219V) protease at 4 °C overnight. Digested Rab5Q80L-$His_{10}$ and the protease were separated with ion exchange on Mono

Q column (GE Healthcare) with MonoQ B buffer (50 mM Tris-HCl pH 8.8, 1 mM DTT, 1 M NaCl). Next, the buffer was exchanged to Rab buffer (20 mM HEPES-KOH pH 7.4, 150 mM KOAc, 1 mM MgCl, 1 mM DTT) and Rab5Q80L-$His_{10}$ was loaded with GTP in presence of 5 mM EDTA and 1 mM GTP for 4 h at room temperature. The reaction was finally quenched with 10 mM $MgCl_2$ and the buffer was exchanged back to Rab buffer using PD-10 columns. Aliquoted samples were then snap frozen in liquid nitrogen and stored at -80 °C.

**Rabex5:Rabaptin5.** The Rabex5:Rabaptin5 GEF complex was expressed in HighFive insect cell culture from bicistronic pFL bacmid expression cassette for 3 days. Rabex5 was cloned downstream of p10 promoter as a (linker)-TEV-$Gly_4$-Rabex5 fusion and TwinStrep-(linker)-TEV-Rabaptin5 expression was under PH promoter control. The harvested cell pellets were resuspended in StrepTrap buffer (50 mM Tris-HCl pH 8.0, 150 mM NaCl, 0.5 mM EDTA, 5 mM 2-mercaptoethanol), supplemented with cOmplete EDTA-free protease inhibitor cocktail, 2.5 U/mL DNase I and 1 mM PMSF. The cells were lysed by douncing and 10 bursts of tip sonication for 1 s with 3 s pause in between. We ultracentrifuged the lysate at 150,000 rcf for 30 min and loaded the supernatant on StrepTrap HP affinity column. The purified complexes were eluted with 2.5 mM D-desthiobiotin and digested with TEV(S219V) protease. Next, the samples were concentrated on Amicon Ultra-2 cassettes and run on Superdex 200 gel filtration column to isolate the digested GEF complexes. The pooled fractions were further concentrated, aliquoted and snap frozen in liquid nitrogen for -80 °C storage. The same protocol was also used for **ΔRabex5**, **ΔRabex5:Rabaptin5** and **Rabex5:$\Delta_{RBD}$Rabaptin5** purification. Importantly, purification of full length Rabex5 resulted in aggregation when not in complex with Rabaptin5. For the fluorescently labeled Rabex5:sCy5-Rabaptin5 complex, Rabex5 was expressed without an N-terminal linker as fluorescent labeling disrupted the Rabex5 catalytic activity. The sortagging reaction was performed after Rabaptin5 digestion with TEV protease.

**RabGAP-5.** Sf9 insect cells were used to express TwinStrep-(linker)-3C-RabGAP-5 construct for 72 h at 27 °C. We used StrepTrap buffer (50 mM Tris-HCl pH 8.0, 150 mM NaCl, 0.5 mM EDTA, 5 mM 2-mercaptoethanol), supplemented with cOmplete EDTA-free protease inhibitor cocktail to resuspend the cell pellets. The cells were lysed by douncing and 1 % IGEPAL CA-630 detergent solubilization for 10 min on ice. Later, the lysate was ultracentrifuged at 150,000 rcf for 30 min at 4 °C. The supernatant was then loaded on StrepTrap HP affinity column and bound proteins were eluted with 2.5 mM D-desthiobiotin. The pooled fractions were digested with HRV 3C protease overnight and the samples were run on Superdex 200 size exclusion column to separate the RabGAP-5 and 3C protease. We finally pooled the RabGAP-5 fractions, which were snap frozen in liquid nitrogen and stored at -80 °C.

Protein labeling

After purification of transgenic proteins and subsequent digestion with TEV(S219V) protease, the proteins were labeled N-terminally with Sortagging method (*4*) using calcium-independent mutant of *S. aureus* sortase A enzyme (SrtA7M) (*5*). First, lyophillized synthetic CLPETGG peptide (Biomatik) was dissolved in Vesicle buffer (20 mM HEPES-KOH pH 7.4, 150 mM KOAc) with 0.2 M TCEP to 50 mM final concentration. In the next step, maleimide-conjugated synthetic fluorescent dye (sulfo-Cy5-maleimide [sCy5-maleimide] or CF488A-maleimide) in DMSO was added to the peptide in 3-times molar excess and left to react at room temperature overnight, protected from ambient light. We quenched the peptide labeling reaction with 1.5 M 2-mercaptoethanol. Labeled peptides were frozen in liquid nitrogen and stored at -80 °C. In the sortagging reaction, the purified protein was mixed with 10 µM SrtA7M and 0.3 mM labeled peptide. The reaction was then incubated for at least 5 hours at 4 °C in dark. Unreacted peptide, dye and SrtA7M were finally removed from the protein sample with size exclusion chromatography.

Coverslip treatment and reaction chamber immobilization

We used 24×50 mm coverslips (no. 1.5H, Marienfeld) in our fluorescence microscopy assays. First, coverslips were cleaned by 1 h incubation in piranha solution (1:3 volume ratio of 30 % $H_2O_2$, Sigma Aldrich : 98 % $H_2SO_4$, Merck) and extensive washing in Milli-Q grade water. The cleaned coverslips were stored in Milli-Q water for up to two weeks. Immediately before use, the coverslips were dried and further cleaned in Zepto B (Diener electronic) plasma oven for 10 min at 30 W under 1 NL/h air flow. We immobilized the microscopy reaction chambers by attaching a cut PCR tube on the cleaned coverslip glass using ultraviolet glue (Norland optical adhesive 63) under 365 nm UV light for 5-10 min. The attached reaction chambers were then ready for supported lipid bilayer preparation.

Supported lipid bilayer preparation

To mimic the intracellular membranes, we prepared supported lipid bilayers (SLBs) on high precision microscope slides (no. 1.5H). The SLBs are made by mixing synthetic lipids with intended composition in chloroform at 1 mM concentration per 0.5 mL volume. In this study, we used 79.9 % DOPC, 20 % DOPS, 0.1 % DMPE-PEG2000 and varying amounts of DOGS-NTA[$Ni^{2+}$] lipids (Avanti Polar Lipids). For the experiments with DOGS-NTA, we decreased the ratio of DOPC proportionally. Next, the lipid mixture was dried under Nitrogen flow in a glass vial and kept under vacuum for at least 1 hour. The dried lipids were hydrated in Vesicle buffer (20 mM HEPES-KOH pH 7.4, 150 mM KOAc) by vortexing to produce multilamellar vesicles (MLVs). In the following step, we prepared small unilamellar vesicles (SUVs) by passing the MLV solution over five freeze-thaw cycles and extrusion through 100 nm polycarbonate membrane 21-times (LiposoFast, Avestin). The produced SUVs were stored at 4 °C for up to 5 days. Similarly, we used DOGS-NTA SUVs within 2 days since production. Finally, the SLB was formed in the immobilized plastic reaction chambers on a clean glass surface by inducing fusion of the SUVs with 3.33 mM $CaCl_2$. The SLBs were left to form for at least 45 min at 37 °C. Later, the unfused vesicles were washed away with Vesicle buffer, Milli-Q water and reaction buffer.

TIRF microscopy

TIRF microscopy experiments were done on (i) Zeiss Axio Observer.Z1 inverted microscope with Visitron iLas2 illumination module (GATACA), Zeiss Deffinite Focus 2 and Plan-APOCHROMAT 63x/NA 1.46 immersion objective and (ii) inverted Olympus IX83 with Cell^TIRF system and Olympus Uapo N 100x/NA 1.49 oil objective. Imaging on the Zeiss system was performed on two Photometrics Evolve-EM 512 D EMCCD cameras. Conversely, the Olympus stand was equipped with water-cooed Hamamatsu C9100-13 EMCCD camera. On both systems, the imaging was done with 200 EM camera gain and 30 ms exposure time with varying laser intensities unless stated otherwise.

Rab5 activation reconstitution assays on SLB

For the *in vitro* reconstitution assays on the SLBs, the purified and fluorescently labeled protein were incubated in the immobilized reaction chambers with glass supported membranes in Rab reaction buffer (20 mM HEPES-KOH pH 7.4, 150 mM KOAc, 1 mM $MgCl_2$, 1 mM DTT, 50 μM GDP), supplemented with 0.5 mM GTP, in 30 μL volume. We used surface-sensitive TIRF microscopy to specifically visualize membrane binding of labeled components. For that, we set the focal plane at the SLB with < 100 nm penetration depth of the evanescent excitation field. We recorded the fluorescence signal in 30 s intervals for at least 10 min after equilibration of the reconstituted system to obtain stable baseline intensity. Then, we injected GEF in 20 uL Rab reaction buffer and mixed the contents of the reaction chamber to initiate the nucleotide exchange. We continued the recording in 30 s intervals until the fluorescence signal reached steady state. For the hysteresis assay, we first injected 10 μL 80 nM Rabex5:Rabaptin5 and allowed the activation reaction to plateau. We induced switching back to the basal state by addition of 10 μL RabGAP-5 to a final concentration of 500 or 2000 μM. In cases where no increase in signal was detected after at least 150 min post-induction, the recording was stopped and the activation delay time was deemed to be >150 min. Additionally, we recorded the camera noise by closing the microscope shutter and collecting the detector readout.

Microscopy data analysis

The collected microscopy data was analyzed using Fiji ImageJ 1.52i package. The membrane localization of the selected fluorescently labeled component was determined by measuring the mean fluorescence intensity at the SLB and subtracting the recorded camera noise value for each recorded frame. The point of GEF addition was set as t = 0. To obtain the time of inflection $T_i$ and maximum growth rate $k_{max}$, we fitted a Gompertz function (*6*) to the subtracted fluorescence intensity values between the GEF addition and the onset of steady state:

$$I(t) = B + (A - B)\exp\left(-\exp\left(-e \cdot k_{max}(t - T_i)\right)\right)$$

where $I(t)$ is the measured fluorescence intensity at time $t$, $A$ and $B$ are the upper and lower fit asymptotes, respectively, $k_{max}$ is the relative maximum signal growth rate and $T_i$ is the temporal delay to reach the $k_{max}$ after the GEF addition. For the reactions where no signal increase was observed after 150 min post-GEF addition, the activation delay $T_i$ was taken to be >150 min and the $k_{max}$ was set to 0. To normalize the data, we divided the fluorescence intensities by the upper fitted asymptote $A$. The traces that did not result in collective activation were normalized by dividing the signal intensities with the mean of upper asymptotes $A$ at the corresponding microscope setup. This workflow is also summarized in Fig. S4. The normalized fluorescence intensities were used to group independent replicates. This way, we obtained the mean intensity traces and standard deviations SD for selected conditions. Similarly, the fold change in fluorescence signal was calculated by taking the upper asymptote value $A$ and dividing it with the mean value of the baseline signal before the GEF injection.

Single particle tracking

For single particle tracking, we used CF488A-Rab5:GDI, supplemented with small mamounts of sCy5-Rab5:GDI. The particle diffusion of sCy5-Rab5 on the SLB was captured using the Zeiss Axio Observer.Z1 inverted microscope with Visitron iLas2 illumination module. We used 100 % 640 nm laser power, 30 ms exposure time, 300 EMCCD camera gain and 100 ms acquisition interval. To capture single particles landing on the SLB before nucleotide exchange, we incubated the glass supported membrane with 500 nM CF488A-Rab5:GDI, 2 μM GDI, 50 μM GDP, 500 μM GTP and ca. 1 nM sCy5:GDI. To limit the effects of photobleaching, we also included an oxygen scavenging system with 60 mM D-glucose, 0.1 mg/ml glucose oxidase (SERVA), 0.32 mg/ml catalase (Sigma Aldrich) and 2 mM Trolox (Sigma Aldrich). We built the particle trajectories using the TrackMate ImageJ plugin v.3.7.0 (*7*). We used simple LAP tracker with 0.7 μm particle diameter, 15 threshold value with median filter, signal-to-noise ratio > 0.6, 2 μm maximum linking distance and up to 2 frame gap with 3 μm closing distance to account for fluorophore blinking. To image particles after the nucleotide exchange with GTP or GMP-PNP, we first triggered the Rab5 activation by injecting 200 nM Rabex5:Rabaptin5 into 500 nM CF488A-Rab5:GDI, 2 μM GDI, 50 μM GDP, 500 μM GTP/GMP-PNP and cca. 50 fM sCy5:GDI mixture. We followed the progression of collective activation by measuring the CF488A-Rab5 fluorescence intensity at the SLB. When the reaction reached steady state, we supplemented the sample with fresh oxygen scavengers and imaged sCy5-Rab5 particles with 100 % 640 nm laser power, 30 ms exposure, 300 EM gain and 100 ms acquisition interval. The TrackMate trajectories were built with simple LAP tracker, 0.7 μm particle diameter, 15 threshold value with median filter, signal-to-noise ratio > 0.6, 1 μm maximum linking distance and at most 2 frame gap with 1.5 μm closing distance. For GMP-PNP-bound sCy5-Rab5, ≤ 0.8 μm and ≤ 1.2 μm distances were used to link particles and close up to 2 frame gaps, respectively. We analyzed only trajectories with 3 spots or longer. To calculate the membrane residence lifetimes for sCy5-Rab5[GDP] $\tau^{GDP}$, we fitted a monoexponential function to the trajectory duration histogram:

$$N(t) = A \cdot e^{-t/\tau} + B$$

where $N(t)$ is the trajectory count number at a given duration $t$ and $\tau$ is the mean lifetime. Conversely, a two-exponential function was better at fitting the sCy5-Rab5[GTP/GMP-PNP] trajectory histogram. This gave us $\tau_1^{GTP/GMP-PNP}$ and $\tau_2^{GTP/GMP-PNP}$ values.

$$N(t) = A_1 \cdot e^{-t/\tau_1} + A_2 \cdot e^{-t/\tau_2} + B$$

The calculated lifetimes are not corrected for photobleaching and thus represent a lower estimate of the actual membrane residence lifetimes.

Activation wave velocity

Activation waves were observed several minutes after 80 nM Rabex5:Rabaptin5 injection to 500 nM CF488A-Rab5:GDI, 2 µM GDI, 50 µM GDP, 500 µM GTP and 50 nM RabGAP-5. The collected time series were first corrected for uneven illumination profile to track the progression of the wave front. To this end, we prepared glass supported membrane, doped with 0.25 µg/mL fluorescent DiO tracer (Sigma Aldrich), to determine the TIRF illumination profile across the field of view. We analyzed the acquired images using ImageJ 1.52i. First, we generated an illumination profile reference image by gaussian filtering of the DiO-labeled SLB snapshot with ImageJ FFT Bandpass Filter. We then divided the wave time series with the illumination reference to produce corrected images. These images were used to produce a kymograph along a line across the field of view. Finally, we estimated the activation wave velocity from the slope of the kymograph fluorescence profile.

Fluorescence recovery after photobleaching

Fluorescence recovery after photobleaching (FRAP) was used to estimate the protein turnover at the SLB after the GEF-mediated nucleotide exchange. A reaction composed of 500 nM CF488A-Rab5:GDI, 2 µM GDI, 50 µM GDP, 500 µM GTP or GMP-PNP and 80 nM Rabex5:Rabaptin5 was let to reach steady state. We then exposed a central area of roughly 20 x 20 µm to high-power 488 laser for 3-5 ms/px. The FRAP was then monitored in 1 s intervals until the fluorescence intensity reached a new steady state. We determined the fluorescence intensities of a center 3×3 px square to minimize the effects of lateral diffusion in the bleached area. To determine the protein exchange rate, we fitted a monoexponential function to the normalized intensity profile:

$$I(t) = A \cdot e^{-k_{ex}t} + B$$

where $I(t)$ is the normalized fluorescence intensity at time $t$ and $k_{ex}$ is the protein exchange rate. We normalized the fluorescence intensity by first subtracting the camera noise from the bleached 3×3 px square intensity profile. Then, we corrected the fluorescence recovery series for photobleaching during imaging by dividing the subtracted intensities with fluorescence signal outside the bleached area. Finally, the corrected fluorescence profiles were normalized to the mean fluorescence intensity of 10 pre-bleach frames.

Size exclusion chromatography – multi angle light scattering

The oligomeric state of the purified Rabex5:Rabaptin5 sample was analyzed with size exclusion chromatography – multi angle light scattering (SEC-MALS). A 100 uL sample at 1.0 mg/mL in 50 mM Tris-HCl pH 7.5, 150 mM KCl, 5 mM $MgCl_2$, 2 mM TCEP and 10 vol% glycerol was run in duplicate on Superdex 200 Increase 10/300 column at 0.5 mL/min and 35 °C. We used OMNISEC RESOLVE for sample separation and the REVEAL module (Malvern Instruments) for multi angle light scattering and refractive index detection. Prior to the runs, the samples were stored at 6 °C in the autosampler. The recorded data were analyzed using the OMNISEC v10.41 software package.

Guanine nucleotide exchange factor assay

Activity assays of purified GEFs were performed in 384-well plates (black non-binding surface microplate, Corining). First, Rab proteins were loaded with mant-GDP by 1 h incubation at 37 °C in presence of 2 mM EDTA and 20-times molar excess of the labeled nucleotide during purification. The exchange reaction was then quenched with 5 mM $MgCl_2$ and the buffer was exchanged to the reaction buffer using desalting columns. On the day of the experiment, 250 nM Rab5[mant-GDP]:GDI, Rabex5:Rabaptin5 and 500 µM SUVs were added to the microplate and incubated in reaction buffer (20 mM HEPES-KOH pH 7.4, 150 mM KOAc, 1 mM $MgCl_2$, 1 mM DTT) for 10 min. The measurements of mant-GDP fluorescence were performed on Biotek Synergy H1 plate reader (excitation 355 nm, emission 450 nm). After acquiring the baseline fluorescence for 40 min, the GEF exchange reaction was induced by injecting GTP to the wells at 1 mM final concentration. Finally, to determine observed exchange rates $k_{obs}$, the measured time courses were fitted with a monoexponential function.

$$I(t) = A \cdot e^{-k_{obs}\, t} + B$$

Where *I(t)* is the measured fluorescence intensity at time *t*. The catalytic efficiency $k_{cat}/K_m$ was obtained from the slope of a linear fit to $k_{obs}$([GEF]) plot. Where $k_{obs}$ is the observed exchange rate at the given Rabex5:Rabaptin5 molar concentration [GEF] and $k_0$ is the intrinsic Rab5 nucleotide exchange rate.

$$k_{obs} = \frac{k_{cat}}{K_m} \cdot [GEF] + k_0$$

Modelling

See Supplementary Text for detailed description of the model.

**Supplementary Text**

Model of Rab5 positive feedback

(i) Reaction scheme

Our model represents a minimal biochemical network (Fig. 1G) taking into account known molecular interactions (*8–14*). The reactions follow a particular sequence: Rab5[GDP]:GDI dissociates into Rab5[GDP] and GDI (Reaction 1 below). Free Rab5[GDP] can subsequently interact with the Rabex5:Rabaptin5 complex, denoted by RR. This leads to the formation of Rab5[GTP] (Reaction 3 below). Rab5[GTP] positively upregulates its own activation through the complex Rab5[GTP]:RR (Reactions 4 and 5 below). Finally, Rab5[GTP] can revert back to Rab[GDP] (Reaction 2 below). We model all reactions using mass-action kinetics. Except where explicitly stated, we assume that molecule diffusion is sufficiently fast so that the system is well-mixed.

The reaction scheme is given by:

(1) $Rab5[GDP]{:}GDI \underset{k_2}{\overset{k_1}{\rightleftharpoons}} Rab5[GDP] + GDI$

(2) $Rab5[GTP] \overset{k_3}{\to} Rab5[GDP]$

(3) $Rab5[GDP] + RR \overset{k_4}{\to} Rab5[GTP] + RR$

(4) $Rab5[GTP] + RR \underset{k_6}{\overset{k_5}{\rightleftharpoons}} Rab5[GTP]{:}RR]5uentlyd\ experiments. Noted\ are\ mean\ lifetimes\ for\ the\ tracked\ part$[illegible]
$exponential\ decay\ fit. les\ with$

(5) $Rab5[GDP] + Rab5[GTP]{:}RR \overset{k_7}{\to} 2Rab5[GTP] + RR$

The positive feedback is implemented through the interplay of the reactions (4) and (5). The last reaction listed likely represents a number of reactions that occur in sequence. Therefore, $k_7$ should be considered as an *effective parameter* that incorporates events such as Rabex5:Rabaptin5 unbinding from Rab5[GTP] and nucleotide exchange of Rab5 from [GDP] to Rab5[GTP]. At sufficiently high concentrations of Rabex5:Rabaptin5 and Rab5[GDP], the system can switch into a stable, activated state. Such ultrasensitive behavior is a well-known property of biochemical networks containing a positive feedback loop (*15*).

(ii) Parameter fitting

We use measurements from our experiments and the literature to constrain the parameter ranges, see Table S1. There is uncertainty in these measurements, so we searched parameter space allowing one order of magnitude change in the parameters. We allowed the parameters involved in the positive feedback (e.g. $k_7$) to vary over a larger range as there is uncertainty in the specific interaction (see below). Due to the time involved in solving the equations stochastically, we did not perform a rigorous search across the complete parameter space. We focused on varying the positive feedback terms to find a parameter set that replicated the behavior shown in Fig. 1. This parameter set (as shown in Table S1), was kept fixed and used in modelling the system perturbations shown in the later results except where explicitly stated otherwise. We note that our final parameters are not unique and that the observed behavior can be qualitatively replicated with other parameter sets. However, in all cases, positive feedback is necessary to reproduce the experimental results.

(iii) Implementation of positive feedback

We emphasize that our model is conservative with regard to the strength of the positive feedback. In Reaction (5), the positive feedback leads to a relatively small increase in Rab5[GTP] number (one Rab5[GTP] goes to two Rab5[GTP]). Further, we constrain the feedback parameters to not be substantially different from other parameters. More complex reactions can be envisaged that lead to stronger feedback, either in terms of the magnitude of the parameters (*e.g.* larger $k_7$) or the number of Rab5 molecules involved in the positive feedback reaction. Examples include release of autoinhibition (*10*) and oligomerization of the ternary complex (*11*). Such increased positive feedback strength will result in more "step-like" behavior and can also make stochastic effects more pronounced. To reiterate, given the lack of details of the biochemical reactions involved, we have focused on developing a minimal model of the Rab5[GTP] positive feedback. Our modeling results show that such feedback is necessary for the switching behavior and the temporal variability at low GEF concentrations. However, we may well be underestimating the magnitude of the positive feedback.

(iv) Solving the reaction scheme

Even in steady-state, the above reactions are challenging to solve analytically. Given that our interest here is in the temporal response of the Rab5 membrane binding, we revert to numerical methods. We assume that the reactions are not diffusion limited – *i.e.* the system is well-mixed. The above reaction scheme was solved using numerical integration with the ODE solver in SimBiology toolbox (*16*) in MATLAB and with a stochastic ODE solver using the Biosimulator.jl package (*17*) in Julia. All stochastic simulations start with 500 particles of Rab5[GDP]:GDI, and varying concentrations of RR and GDI (based on the experiments). All other reaction species have zero particles at the start of the experiment. The relative magnitude of the stochastic fluctuations depends on the local particle number.

The model, when solved stochastically, captures the following aspects observed experimentally, Fig. 1G-J and Fig. 3E:

- The delay of Rab5 activation at intermediate RR concentrations and the time for Rab5 activation decreases with increasing RR concentration;
- At intermediate RR levels, the Rab5[GTP] concentration remains near background levels until switching rapidly to the "ON" state.
- The change in the relative maximum rate of the Rab5 activation, with varying RR concentration;
- Decreasing activity with increasing GDI activity.

When we solved the model deterministically, we obtain qualitative similar results to the stochastic solution except: (1) we have gradually increasing Rab5[GTP] levels at intermediate RR levels before switching, whereas experimentally this is not seen; and (2) we do not get the temporal variability in the switching times at intermediate concentrations. At high RR levels the stochastic and deterministic solutions are similar as fluctuations are relatively small (Fig. S6).

We cannot discount that the system behavior can be described entirely through a deterministic approach. One method would be to introduce parameter variability (rather than stochastic noise) to recreate the distributed activation times at intermediate RR concentrations. However, given the precisely controlled experimental conditions, protein number stochasticity provides a simple explanation for our results without requiring additional complexity.

<u>Phenomenological model of Rab5 cascade</u>

To explore the parameter space for stochastic switching and to test the interplay between the positive feedback and the initial condition of the system, we constructed a phenomenological model of Rab5 activation. This phenomenological model captures the time-delayed response and relative maximum rate curves (Fig. S8). However, reproducing more complicated interactions, such as changes involving GDI concentration, requires the full model as outlined above.

Our phenomenological model uses a Hill function to implement the positive feedback of membrane bound, GTP-activated Rab5 on its own activation:

$$\frac{d[R]}{dt} = a_0 + a_1 \left(\frac{[GEF]}{100nM}\right) \frac{[R]^n}{[R]^n + K^n} - a_2[R] \qquad [1]$$

$[R]$ represents the membrane bound, GTP-activated population of Rab5. $a_0$ is the basal activation rate, $a_1$ is the positive feedback strength, $K$ is the effective $K_D$ of the positive feedback reaction, and $a_2$ is the Rab5 membrane disassociation rate. In the absence of positive feedback, the steady-state concentration is $a_0/a_2$.

Such a model can be derived as an approximation of the above reaction network with the following assumptions: (1) the slowest reaction is the positive feedback and effectively all other components are near equilibrium; (2) the levels of [Rab5:GTP] are significantly larger than [Rab5:GDP]; and the reaction from [Rab5-GTP:RR] to [Rab5-GTP]+[RR] is negligible. Under these assumptions, $a_0 \sim \frac{[RR]_T\Phi}{G}$ and $a_1 \sim \frac{[RR]_T}{G}$ and *n=2*, where $[RR]_T$ is the total concentration of Rabex5:Rabaptin5, Φ is the total amount of Rab5 (in all forms) and G is the total amount of GDI. Related to the above discussion regarding the strength of the positive feedback, it is straightforward to envisage larger effective Hill coefficient (*e.g.* $n = 4$). However, given uncertainty about the precise mode of feedback we take a conservative approach with *n=2*.

Without loss of generality, we define $t = \tau / a_2$. Substituting into Eq. 1, we obtain

$$\frac{d[R]}{d\tau} = b_0 + b_1 \frac{[R]^n}{[R]^n + K^n} - [R] \qquad [2]$$

where $b_0 = a_0/a_2$ and $b_1 = a_1/a_2$; both $b_0$ and $b_1$ have dimensions of concentration.

This system has only one stable (low activity) state for $b_1 \ll b_0$. At large enough $b_1$ the system also has a stable high activity state. In the case of $b_0=0$, and *n=2*, this corresponds to $b_1 > 2K$. See Lewis *et al.* for a detailed discussion of the system stability (*18*).

Here, we are interested in the time-dependent behavior of this system given the inherent stochasticity of protein-protein interactions. We took two approaches: (i) Gillespie algorithm simulations of Eq. [2]; and (ii) solving the corresponding Fokker-Planck equation for Eq. [2]. Finally, in (iii) we discuss how Eq. [2] can be adapted to give rise to spatial waves of activation.

(i) Simulations

Simulations were performed using a Gillespie algorithm in Matlab. To test our code, we first performed simulations with $b_1=0$. In this case, Eq. [2] can be solved exactly for the stochastic case through a Master Equation approach (*19*). We confirmed that the measured concentration mean and fluctuations were as predicted. Positive feedback was then implemented as a separate term within the propensity function (*20*). Each simulation was run for $2 \times 10^4$ time steps and for each parameter set the simulation was repeated four times (Fig. S8). All results shown are for $n = 2$. We confirmed that qualitatively similar results can be obtained for $n = 4$ (not shown). Simulations always started with zero bound population of Rab5. See Table S2 for simulation parameters.

To implement hysteresis, as shown in Fig. 4E, for each parameter set the system is either started at either $a_0/a_2$ (for OFF to ON) or $\left((a_0 + \mathrm{a}_1 \frac{[\mathrm{GEF}]}{100\mathrm{nM}}\right)/a_2$ (for ON to OFF). The number of particles in the simulation was recorded after 150 minutes.

(ii) Fokker-Planck Equation

Assuming sufficiently large particle number, we can use a Fokker-Planck approximation for the probability of membrane bound GTP-activated Rab5 having a particular concentration $A$, *P(A)*, as (note, here we switch to $[R] = A$ to simplify nomenclature):

$$\partial_t P(A) = \frac{1}{2}\partial_x^2[D(A)P(A)] - \partial_x[v(A)P(A)] \quad [3]$$

Where *D* and *v* represent the effective diffusion and drift terms. See Walczak *et al.* (*19*) for discussion of the derivation of Eq. [3]. We can re-write Eq. [2] in terms of a generalized diffusion-drift equation;

$$D(A) = b_0 + b_1 \frac{A^2}{K^2+A^2} + A \quad [4a]$$

$$V(A) = b_0 + b_1 \frac{A^2}{K^2+A^2} - A \quad [4b]$$

Although difficult to make analytical process, it is straightforward to solve Eqs. [3-4] numerically. We use zero diffusive flux boundary conditions, with initial condition $P(0) = 1$ and $P(x>0)=0$. We solve Eq. 3 using the Matlab solver *pdepe*. There is no source term, and hence the total probability at each time point is one (used as a check to ensure the solver worked correctly). We first confirmed that for the case $n=0$ – where Eqs. [4a-b] reduce to the case of Brownian motion – our solution is consistent with the known Gaussian solution to Brownian motion, with mean and variance equal to $b_0 + b_1$.

We next explored $n>0$. The critical point at which the distribution becomes bimodal depends sensitively on *n*; see (*19*) for detailed discussion. From the distribution of *P(A)* we can predict the switching time for the system given specific parameters and starting from the OFF state. In particular, we can see how the switching time depends on the positive feedback (Fig. S9).

We can test dependency of switching on $b_0$. Insufficient basal production and the system never reaches an excited state. However, if $b_0$ is too large, the system effectively loses any switching behavior as the difference in concentration between "on" and "off" states is small. We define the switch score S as:

$$S = (b_2/b_0)[A]_{ss} - 1 \quad [5]$$

When there is no positive feedback, *S=0.* We define a "good" switch as having *S > 10. i.e.* the final steady-state is at least ten times larger than the stable state ($[A]_{ss} = b_2/b_0$) in the absence of positive feedback. These results are shown in Figure 3 for *K=100* for different parameter values of $b_0$ and $b_1$. As *K* is changed, the general shape of the regime varies as the number fluctuations change. In Fig. S9, we show as an example the parameter space for K=30.

(iii) Spatial activation waves

To incorporate spatial propagation of membrane bound GTP-activated Rab5, we incorporate membrane-bound diffusion of Rab5 into Eq. [1].

$$\frac{\partial A}{\partial t} = D\frac{\partial^2 A}{\partial x^2} + a_0 + a_1\frac{\left(\frac{A}{A_0}\right)^n}{1+\left(\frac{A}{A_0}\right)^n} - a_2 A \quad [6]$$

Definitions:
$A$ = concentration of GTP-activated Rab4 on surface $[\mu m^{-2}]$
D = diffusion coefficient of protein on surface $[\mu m^2\ s^{-1}]$
$a_2$= membrane disassociation rate of surface-bound protein [$s^{-1}$]
$a_0$= spontaneous membrane binding rate $[\mu m^2\ s^{-1}]$
$a_1$ = positive feedback strength $[\mu m^2\ s^{-1}]$
n = Hill coefficient on positive feedback
$A_0$= threshold response of positive feedback $[\mu m^{-2}]$

This can be rewritten in terms of the dimensionless variables $A = A_0\phi, t = \tau/a_2, x = u\sqrt{\frac{D}{a_2}}, \widetilde{a_1} = \frac{a_0}{a_1\rho_0}, \alpha = \frac{a_1}{a_2 A_0}$, where $\phi, \tau, u$ are dimensionless concentration, time and position respectively (*21*).

$$\frac{\partial \phi}{\partial \tau} = \frac{\partial^2 \phi}{\partial u^2} - \phi + \widetilde{a_1} + \alpha\frac{(\phi)^n}{1+(\phi)^n} \quad [7]$$

The system is reduced to three parameters. We consider *n =2*; although larger *n* changes the system response, the qualitative behavior is similar. Here, we typically consider $\widetilde{a_1} \ll \lambda$. We explored the propagation of the Rab5 activation once it is activated at a specific point (Fig. 4H). Stochasticity is less relevant in this regime, as we consider the system behavior after Rab5 activation has already been triggered.

Under what conditions does Eq. 7 support waves of activation? Interestingly, whether waves with constant speed (as seen experimentally, Fig. 4E) exist is largely independent of the value of *D* itself, though if the waves exist $D$ determines the wave speed. Considering $a_0=0$ for now, the system is described by one dimensionless parameter:

$$\alpha = \frac{a_1}{a_2 A_0} \qquad [8]$$

Though precise details depend on the values of $a_0$ and $n$, stable waves with constant velocity typically exist if $\alpha \geq 2$ (*21*). Unsurprisingly, stronger positive feedback results in more stable and faster waves; *i.e.* the wave can propagate reliably. Increasing the disassociation rate ($a_2$), and/or the threshold concentration for the positive feedback ($A_0$), decreases wave propagation. This makes intuitive sense: if protein disassociates rapidly, then the wave will struggle to propagate; and if the threshold for the positive feedback is high then propagation of the wave needs high local concentrations of protein, reducing the chance of it spreading. Of course, more complex wave-like behavior is possible. Our objective here is not to precisely model the wave propagation, but to show that our simple model is able to qualitatively replicate the observed wave dynamics with a minimal parameter set.

For waves of constant speed, the wave speed in one-dimension is given by $c \approx \sqrt{\frac{Da_1}{A_0}}\left(1 - \frac{3}{2}\frac{A_0}{a_1} a_2\right)$ (*21*), where $a_0 = 0$. Larger $D$ or stronger positive feedback propagate the wave more rapidly. Increasing the threshold for positive feedback reduces the wave speed. Including $a_0>0$ does not change the fundamental behavior of the system for small $a_0$, but it does shift the allowable parameter values. At high $a_0$ the membrane-bound lifetime is largely independent of position as local spontaneous activation is much more likely and not via an incoming wave of activation.

For the results shown in Fig. 4, we assume that Rab5 activation is triggered from a point source in two-dimensions. Therefore, Eq. 6 is adapted so that the diffusive term is solved in plane polar coordinates $\left(\frac{1}{r}\frac{\partial}{\partial r} + \frac{\partial^2}{\partial r^2}\right)\phi$. With this change, we solved Eq. 7 using *pdepe* solver in Matlab.

In the experiments, by tuning the [GAP] concentration we are effectively altering the protein disassociation rate. Increased [GAP] corresponds to effectively increasing $a_2$. So, larger [GAP] concentrations steadily slow the wave, until it abruptly disappears. Without the presence of the GAP, it is likely that the trigger is too rapid to observe in our experimental setup (Figs. 1-3).

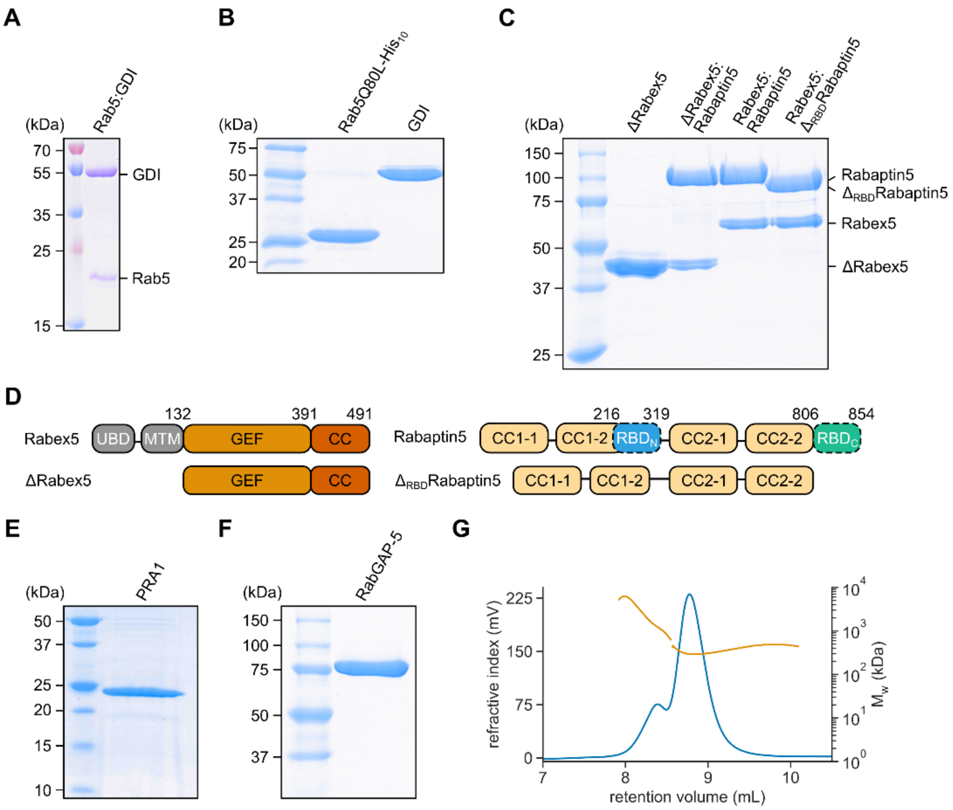


| | Peak 1 | Peak 2 | Peak 3 |
|---|---|---|---|
| retention volume (mL) | 7.87 ± 0.01 | 8.54 ± 0.01 | 8.78 ± 0.01 |
| Mw (kDa) | 73740 ± 8296 | 1990 ± 73 | 323 ± 4 |
| Frac. of sample (%) | 0.8 ± 0.4 | 21.8 ± 1.4 | 77.4 ± 1.8 |

**Fig. S1. Purified protein components used in this study.**

**(A)** Purified *X. laevis* Rab5:GDI complex. The expected molecular weights were 23.7 and 50.8 kDa for Rab5 and RabGDI, respectively. **(B)** Rab5Q80L-His$_{10}$ and RabGDI. The expected molecular weights were 24.7 kDa for Rab5Q80L-His$_{10}$ and 50.8 kDa for RabGDI, respectively. **(C)** The purified GEF complexes. Expected molecular weights for monomeric components: 42.0 kDa, 57.1 kDa, 81.1 and 99.2 kDa for ΔRabex5, full length Rabex5, Δ$_{RBD}$Rabaptin5 and Rabaptin5, respectively. **(D)** Schematic representations of Rabex5 and Rabaptin5 deletion mutants used in Fig. 2B. UBD: ubiquitin binding domain; MTM: membrane targeting motif; GEF: GEF domain; CC: coiled-coil region; RBD: Rab5 binding domain. **(E)** Purified PRA1 in 0.1 % Triton X-100. The PRA1 had expected molecular weight of 21.3 kDa. **(F)** RabGAP-5 (SGSM3) with expected molecular weight 86.1 kDa. **(G)** SEC-MALS characterization of purified Rabex5:Rabaptin5 complex. The sample was run in duplicate and produced three peaks. The first peak (< 1 % of total sample) are protein aggregates, the second peak represents higher-order oligomers with average molecular weight 1990 kDa (22 %) and the third peak is 2:2 Rabex5:Rabaptin5 heterodimer with average molecular weight 323 kDa. The expected molecular weight of a 2:2 complex is 313 kDa. The second peak with corresponding apparent molecular weight of 1190 kDa could represent a hexamer of 2:2 heterodimer GEF complexes.

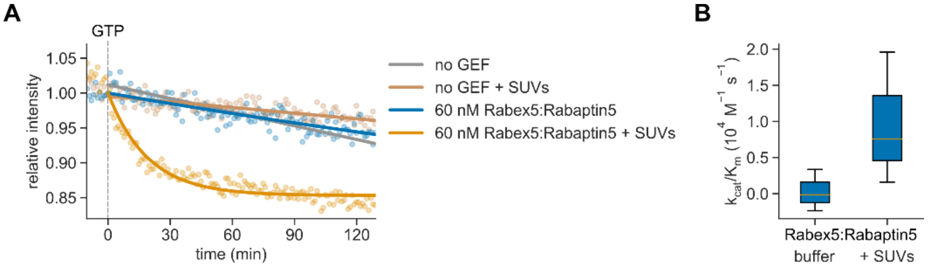


**Fig. S2. Membranes are necessary for Rab5:GDI activation.**

**(A)** Kinetic traces of Rabex5:Rabaptin5 activity assay on Rab5:GDI. The 250 nM mant-GDP loaded Rab5:GDI complexes were incubated with 60 nM Rabex5:Rabaptin5 in reaction wells in presence or absence of SUVs. The Rab5 activation reaction was monitored by relative change in intrinsic mant-GDP fluorescence after nucleotide exchange with non-labeled GTP, which was injected at t = 0. The GEF-mediated nucleotide exchange proceeds only in presence of 500 nM SUVs, signified by marked exponential decrease in signal intensity. Points are means from three independent experiments; solid lines are linear and monoexponential fits for buffer control and experiments with Rabex5:Rabaptin5, respectively. **(B)** Catalytic efficiency of the purified GEF complex on Rab5:GDI in reaction buffer and in reaction buffer, supplemented with SUVs as reaction substrates (n = 3). The determined Rabex5:Rabaptin5 catalytic efficiency in presence of membranes was $1.0 \pm 0.7 \cdot 10^4$ $M^{-1}$ $s^{-1}$.

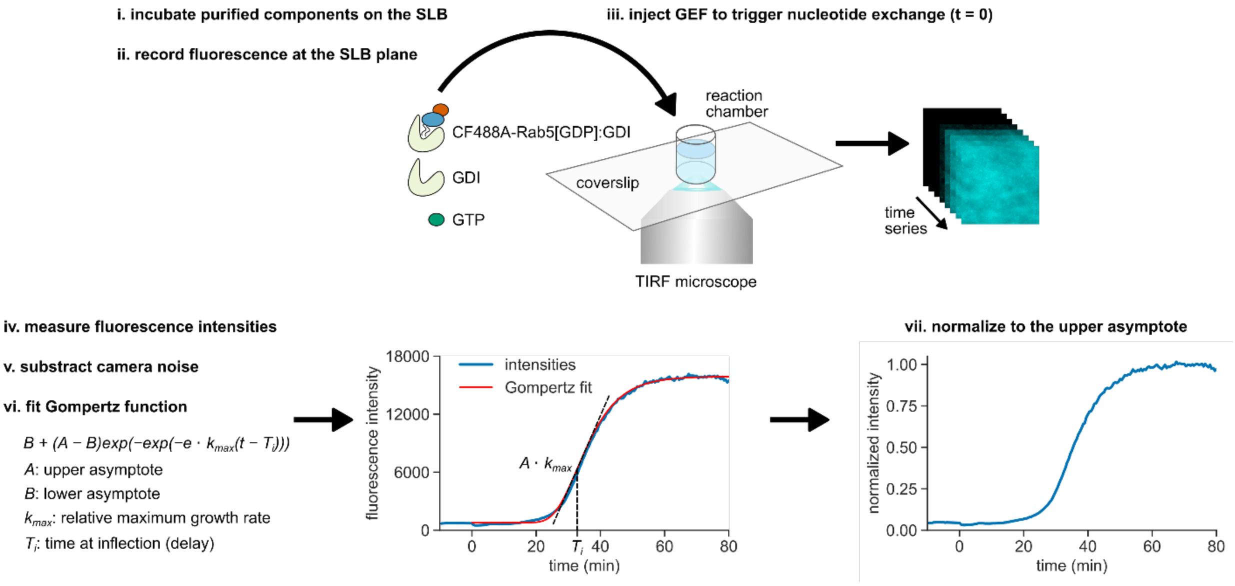


**Fig. S3. Workflow of Rab5:GDI activation analysis.**

First, we prepared coverslip-supported lipid bilayer in immobilized plastic reaction chambers, where we incubated purified protein components (i). The basic reaction included fluorescently labeled Rab5:GDI, free GDI and GTP. We used surface-sensitive TIRF microscopy to specifically observe membrane binding events of labeled components from the solution (ii). After the system equilibrated, we induced the Rab5 nucleotide exchange by the GEF complex addition (iii). The course of Rab5 membrane recruitment was captured as a time series, which was used to measure mean frame fluorescence intensities (iv). After subtraction of camera noise (v), we fitted a sigmoidal Gompertz function to the data (vi). This way, we obtained the relative maximal activation rate $k_{max}$ and the temporal delay until this point of inflection $T_i$. Finally, in order to compare individual runs, we normalized the measured arbitrary fluorescence intensities to the upper asymptote of the Gompertz fit $A$ (vii).

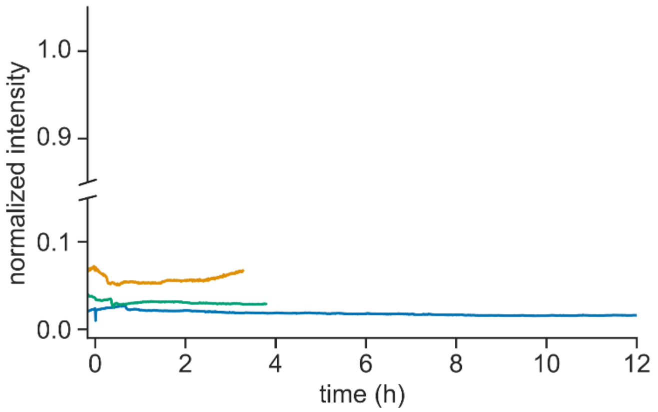


**Fig. S4. Low amounts of Rabex5:Rabaptin5 fail to trigger collective switching even after 12 hours.**

Prolonged kinetic traces of Rab5:GDI switching with 8 nM Rabex5:Rabaptin5. CF488A-Rab5[GDP]:GDI was incubated with GDI, GTP and 10-times lower amount of GDP in an immobilized reaction chamber above glass supported membrane. After 10 min, 8 nM Rabex5:Rabaptin5 was injected and CF488A-Rab5 fluorescence signal was tracked with TIRF microscopy at the SLB focal plane in 30 s intervals for 12 hours. Final concentrations were: 500 nM CF488A-Rab5:GDI, 2 µM GDI, 0.5 mM GTP, 0.05 mM GDP and 8 nM Rabex5:Rabaptin5.

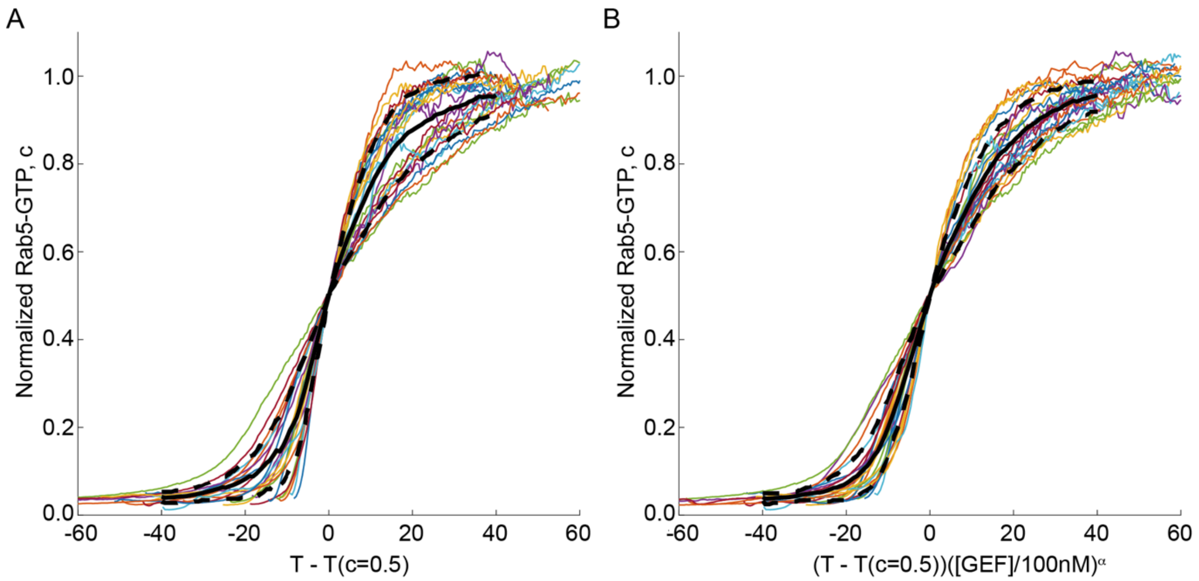


**Fig. S5. Collective Rab5 activation switching follows similar sigmoidal time profile.**
**(A)** Every experimental profile for Rab5 activation with GEF concentrations from 40 nM to 200 nM (n=24). Time axis is centered at 0 for each profile, defined by when each profile passes through 0.5 of final (normalized) signal. Black line represents mean profile and dashed black lines ± 1 SD. **(B)** Same as (A), but with scaled time axis ($\alpha$ = 0.39). Though such scaling decreases variability, the curves do not overlap precisely.

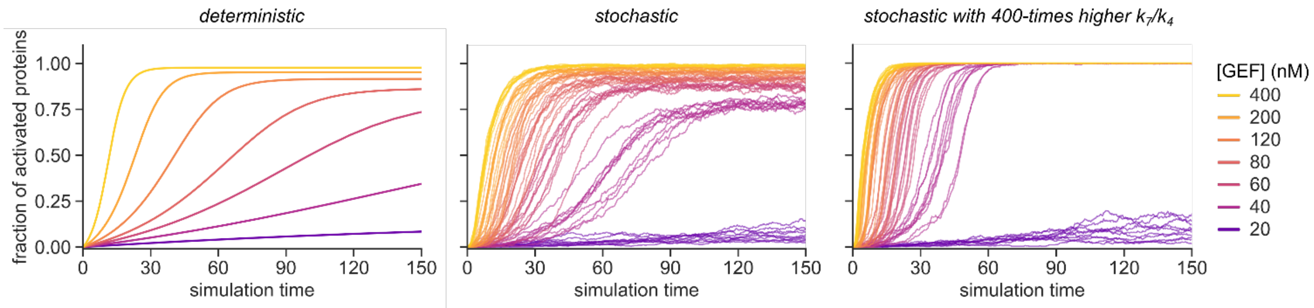


**Fig. S6. Stochasticity is crucial component of the modeled protein network.**

Using our model of Rab5 positive feedback, we constructed a set of ODEs and solved them either deterministically (left) or stochastically (middle and right). Only by running the stochastic simulations, we could observe the characteristic delays in activation and sigmoidal reaction profiles near the critical GEF concentration (see Fig. 1C). At high GEF concentrations, the stochastic and deterministic solutions are comparable, in agreement with diminished stochastic effects. Increasing the $k_7/k_4$ ratio (i.e. greater positive feedback effect) reduces the temporal delay and increases rates of collective switching. Under these conditions, the critical GEF concentration is maintained above 20 nM. The stochastic simulations were run 50-times per condition. Depicted are 10 random runs per given GEF concertation. Simulation parameters are listed in Table S1.

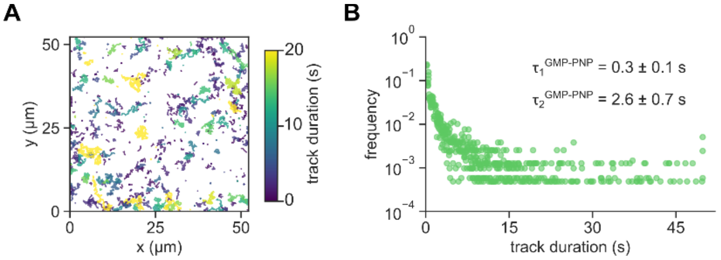


**Fig. S7. GMP-PNP-bound Rab5 single particle trajectories.**

**(A)** Single particle trajectories for activated sCy5-Rab5[GMP-PNP] in steady state. Before acquisition, CF488A-Rab5:GDI, doped with sCy5-Rab5:GDI, was activated with 200 nM Rabex5:Rabaptin5 in presence of GMP-PNP. Shown are 1000 trajectories, which are color-coded for their duration. **(B)** Frequency histogram for sCy5-Rab5[GMP-PNP] trajectory duration from three independent experiments. Noted are mean lifetimes ± SD (n = 3) for the tracked particles according to a two-exponential decay fit.

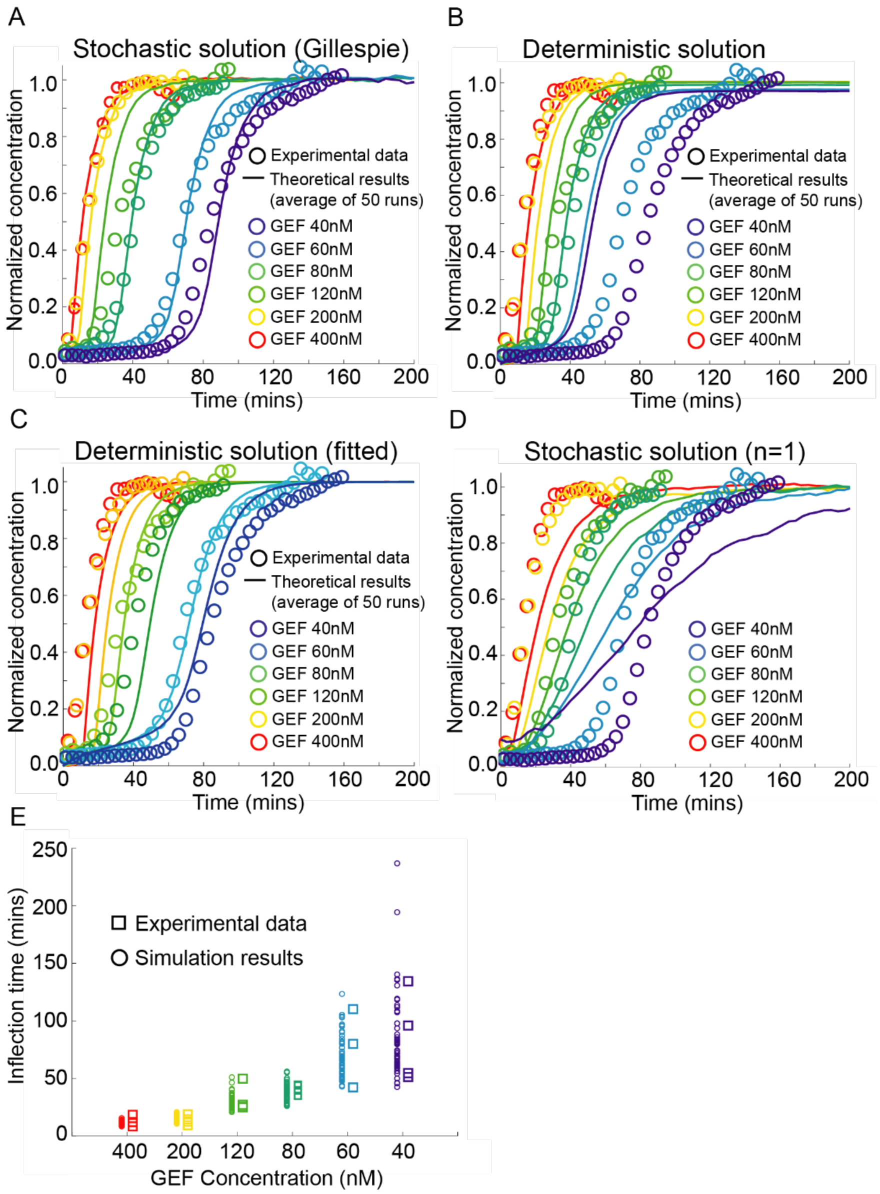


**Fig. S8. Solving the phenomenological model stochastically can qualitatively reproduce GEF titration results.**

**(A-D)** Fitting of phenomenological model (Eq. 1) to experimental data. Theory curves represent 50 simulations per condition. For each condition, the four experimental curves are adjusted such that the time at which concentration = 0.5 is the same, then they are averaged. (A) Eq. 1 solved stochastically for n=2. (B) Eq. 1 solved deterministically using same parameters as (A). (C) Eq. 1 solved deterministically where all parameters as (A) except $a_1$ which is scaled to find best fit to data. (D) Eq. 1 solved stochastically for n=1, with all other parameters kept constant. **(E)** Time for concentration to reach 0.5 of final concentration in both experiments (circles) and simulations (squares, taking individual simulations that form the mean shown in A).

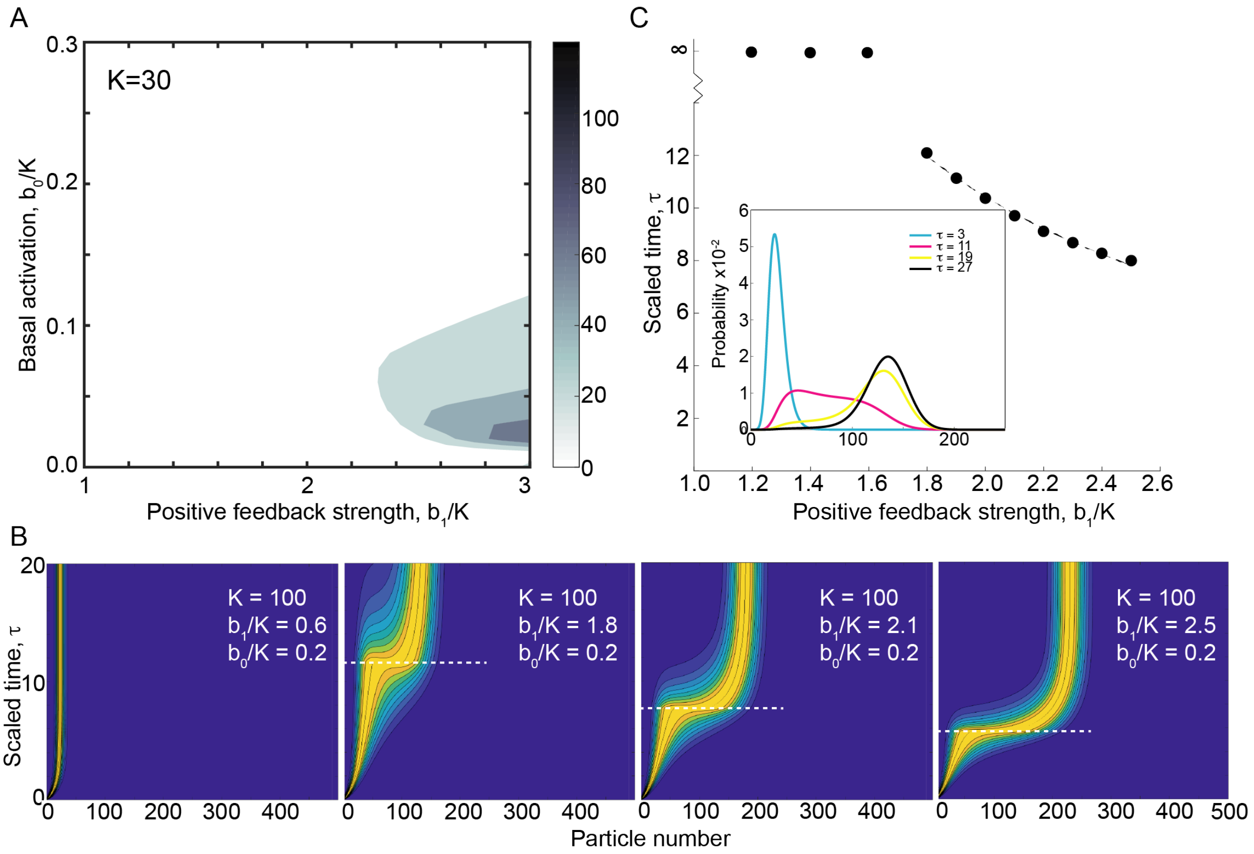


**Fig. S9. Fokker-Planck solution of phenomenological model.**

**(A)** As Fig. 3C except K = 30. $b_0/K$ represents the basal activation rate. $b_1/K$ represents the positive feedback strength. **(B)** Probability density maps of Fokker-Planck solution to phenomenological model for different levels of positive feedback (shown in inset to each panel). Dash white lines denote switching time. y-axis is scaled time (see discussion around Eq. 4). x-axis represent particle number. At small $b_1/K$, the system cannot switch into an ON state (left panel). Blue represents zero probability and bright yellow represents one. **(C)** Switching time from OFF to ON as positive feedback is tuned. For $b_1/K < 1.75$ no switching is observed. Dashed line is fit to $c_1 \left(\frac{b_1}{K}\right)^{-c_2} + \tau_\infty$, where $c_2 = 1.35$ and $\tau_\infty = 0.4$, where $\tau_\infty$ corresponds to switching time for system with $b_1 \gg K$ for given $b_0$. Inset shows probability distribution at different times for K=100, $b_0/K = 0.2$, $b_1/K = 1.8$. Magenta curve corresponds to profile along white dashed line in the second panel of (B).

**Table S1. Reaction parameters used in the model of Rab5 positive feedback.**

| Parameter | Value | Notes |
|---|---|---|
| $k_1$ | $1.2 \times 10^{-1}\ s^{-1}$ | Fitted parameter from estimated $K_d = k_1/k_2 = 6\ \mu M$ (*22*). |
| $k_2$ | $2.25 \times 10^{5}\ M^{-1}s^{-1}$ | Fitted parameter from estimated $K_d = k_1/k_2 = 6\ \mu M$ (*22*). |
| $k_3$ | $5.25 \times 10^{-4}\ s^{-1}$ | Fitted parameter from $k_3 = 0.0006\ s^{-1}$ (*11*, *23*). |
| $k_4$ | $5 \times 10^{4}\ M^{-1}s^{-1}$ | Fitted parameter from $k_4 = 2.5 \cdot 10^4\ M^{-1}\ s^{-1}$ (*8*). |
| $k_5$ | $2.875 \times 10^{4}\ M^{-1}s^{-1}$ | Fitted parameter from estimated $K_d = k_5/k_6 = 50\ \mu M$. |
| $k_6$ | $8.0 \times 10^{-2}\ s^{-1}$ | Fitted parameter from estimated $K_d = k_5/k_6 = 50\ \mu M$. |
| $k_7$ | $10 \times 10^{7} M^{-1}s^{-1}$ | Fitted parameter from $k_7 = 7.5 \cdot 10^4\ M^{-1}\ s^{-1}$ (*8*). |

**Table S2. Reaction parameters used in Gillespie simulations of phenomenological model.**

| Parameter | Value |
|---|---|
| $a_0$ | *0.023* |
| $a_1$ | *0.95* |
| $a_2$ | *0.0018* |
| $n$ | *2* |
| $A_0$ | *100* |

These parameters are used for all data sets except for 40nM GEF concentration, where we take $a_1=1.25$.

**Table S3. Sequences of purified components used in this study.**

| Name | Sequence | Notes |
|---|---|---|
| / | CLPETGG | N-terminal sortagging synthetic peptide (*4*) (Biomatik). The N-terminal Cys was tagged with maleimide-conjugated fluorescent dye. |
| GDI | GADEEYDVIVLGTGLTECILSGIMSVNGKK VLHMDRNPYYGGESSSITPLEELYKRFDMA DGPPESMGRGRDWNVDLIPKFLMANGQLVK MLLYTEVTRYLDFKVIEGSFVYKGGKIYKV PSTETEALASNLMGMFEKRRFRKFLVFVAN FDENDPKTFEGVDPMDTNMRDVYKKFDLGQ DVIDFTGHALALYRTDDYLDQPCLETINRI KLYSESLARYGKSPYLYPLYGLGELPQGFA RLSAIYGGTYMLNKSVDEIVMEKGTVVGVK SEGEVARCKQLICDPSYVPDRVHKAGQVIR VICILNHPIKNTNDANSCQIIIPQNQVNRK SDIYVCMISYAHNVAAQGKYIAIVSTTVET AEPEKEIEPALELLEPIEQKFMAISDLYES TEDGTESQIFCSRSYDATTHFETTCNDIKD IYKRMTGTDFDFENMKRKQNDVFGEDEQ | *X. laevis* RabGDI isoform alpha was expressed as $His_6$-(linker)-TEV-GDI fusion and digested with TEV(S219V) protease. |
| PRA1 | GGGGGAGKNGDDFSDVAEEGPGGILNKMFP KMITQTAAKDWINRRRAHIRPWRNFVDQRR FSRPPNFGELCKRMTRNVEHFQSNYIFIFL GLILYCIITSPMLLIALAVFFGGCYIIYLR TLESKMVLFGRELSTANQYGLAGAVSFPFF WLAGAGAAVFWVIGATLVVIGSHASFHEIE GEVEELQMEPV | *X. laevis* PRA1 was expressed as TwinStrep-(linker)-TEV-$Gly_4$-PRA1 fusion and digested with TEV(S219V) protease. Contains four predicted transmembrane helices. |
| Rab5 | GGGGGANRGGATRPNGPNAGNKICQFKLVL LGESAVGKSSLVLRFVKGQFHEFQESTIGA AFLTQTVCLDDTTVKFEIWDTAGQERYHSL APMYYRGAQAAIVVYDITNEESFARAKNWV KELQRQASPNIVIALSGNKADLSTKRAVDF QEAQAYADDNSLLFMETSAKTSVNVNEIFM AIAKKLPKTEPQAGASNTIRGRGVDLTETA QPTKSQCCSN | *X. laevis* Rab5A was expressed as TwinStrep-(linker)-TEV-$Gly_4$-Rab5A fusion and digested with TEV(S219V) protease. The C-terminal Cys residues were post-translationally geranylgeranylated in insect cells. |

| | | |
|---|---|---|
| Rab5Q80L-His10 | GGGGGANRGGATRPNGPNAGNKICQFKLVLLGESAVGKSSLVLRFVKGQFHEFQESTIGAAFLTQTVCLDDTTVKFEIWDTAGLERYHSLAPMYYRGAQAAIVVYDITNEESFARAKNWVKELQRQASPNIVIALSGNKADLSTKRAVDFQEAQAYADDNSLLFMETSAKTSVNVNEIFMAIAKKLPKTEPQAGASNTIRGRGVDLTETAQPTKSQHHHHHHHHHH | GTPase-deficient Rab5 *X. laevis* Q80L mutant was expressed as TwinStrep-(linker)-TEV-Gly4-Rab5Q80L-His10 fusion and digested with TEV(S219V) protease. The numbering refers to wild-type protein. |
| Rabaptin5 | GAAEPGSVVQPDATLQQRVQELERDNAEFLRTKQLLEQEFNQKRAKFKELYLSKEEDLKHQQAVIQVAQEEIVQLNIKLSQAQAEMENIKAVATVSENTKQEAIDEVKKQWQEEVASHQAIMKETVREYELQFHHRLEQERAQWGQYRESVEREIAELRRRLSEGQQEENLENDMKKAQEDAEKLRSVVMPMEKEIGTLKEKLTEAEEKIKDLEASKMKEMNHYLEAEKSCRTDLEMYVAVLNTQKSVLQEDAEKLRKELHEVCHLLEQERQQHNQLKHTWQKANDQFLESQRLMMQDMKRMEAVLTTEQLRQVEESKKKNQVEEQRTRKRKEKETVQKEEECRKVILEESLPNLKQEELLNSSHSSIHSLDTDIMLHEGDSFNKQEDLFKDGLRRAQSSDSLGASGPLQTKTLGYNNKAKSAGNLDESDFGPLVGADSVSENFDTSSLGSLHMPSGFMLTKDQEKAIKAMTPEQEETASLLSSVTQTLECTYVPPSDYRLVSETEWNLLQKEVQTAGNKLGRRCDMCSNYEKQLQVIQTQEAEIRDQVKKLQTMLRQVNDQLEKTLKDKKDLEDYMKQNTEETSTQISTLTLRINQSETLLADLQQAFHTGKRNIQDQMAVLMHSREQVSEELLRLQRDNESLQGKHSLHVSLQQSEIFNLPETSEELQNLVLKYREDIISVRTATDHLEEKLKAEILFLKEQIQAEQCQKENIEETLQIEIENCKEEMASTSSLQLELDRIKAEREQLEVSLQEKTQELQNLQTLKDSLENQLKKETGSKASLEQLAFEEKNKAQRLQTELDVSEQVQRDFVKLSQMLQVQLERIRQTESLETIRAILNDTKLTDINQLPET | *X. laevis* Rabaptin5 (RABEP1) was expressed as TwinStrep-(linker)-TEV-Rabaptin5 fusion and digested with TEV(S219V) protease. Also stable when expressed alone. |
| Rabex5 | GGGGGSLKTERRGIHVDQSELLCKKGCGYYGNPAWQGFCSKCWREEYQKARQKQIQEDWEFAERLQREEEEAYASSQGAQAGPQSLTFSKFEEKKSNEKTRKVTTVKKFFTASSKSLPKK | *X. laevis* Rabex5 (RABGEF1) was expressed as (linker)-TEV-Gly4-Rabex5 fusion and digested with TEV(S219V) |

| | | |
|---|---|---|
| | DIKEAKSPSPSLSRQFSLETDRVSKDFIEF LKTYQKAGHDVYKLSKIFLEAMHHKRESNI DEQSEFTQDFYQNTADKLQMYWKVSPDKVE KVMDQIERFIMTRLYKHVFCPETTDDEKKD LTVQKRIRALHWVTLQMLCVPVNEDIAEVS DMVVKAITDIIEMDSKRIPRDKLACITRCS KHIFNAIKITKNEPASADDFLPTLIYIVLK ANPPRLQSNIQYITRFCNPSRLMTGEDGYY FTNLCCAVAFIEKLDGQSLNLSEEEFSRYM SGQASPKKQDLENWPEDTCTGVKQMHRNLD LLTQLSKRQEHIVNGAKKLEKDLIDWTDEV TKEVKDIVEKYPLNIKTASQALAALESENV EDDNLPPPLQPQVYAG | protease. Only soluble as Rabex5:Rabaptin5 heterodimer. For the labeled Rabex5:sCy5-Rabaptin5, the Rabex5 N-terminal sequence was MSLKT… to prevent labeling of the Rabex5 moiety. |
| RabGAP-5 | GPSGSYTPSPGGPFSALTASMWPQDILAKY TQKEQTVEQPEFRYDEFGFRVDKEDGAEPN SSKLLGIPLTEDPQQRLRWQAHLEFTHNHD VGDLTWDKIDVTLPHSDKLRSLVLAGIPHS MRPQLWMRLSGALQKKQNSEMTYKDIGRNS SNDDTLAAKQIEKDLLRTMPSNACFSNLQS VGVPRLRRVLRGLAWLFPDIGYCQGTGMVA ACLLLFLEEEDAFWMMAAIVEDLVPVSYFN TTLVGVQTDQRVLRHLIVQYLPRLDKLLQE HDIELSLITLHWFLTAFASVVHIKLLLRIW DFFFYQGSLVLFQTTLGMLKMKEEELIQSE NSASIFNTLSDIPSQIEEADVLLREAMLIS GTLTEVMIEAQRRKHLAYLIADQGQLLNST AAVANLSKIMRRQSQRRKSAITTLLFGDDN FEALKSKNIKQTALVADLREAILQVARHFQ YTDPKNCSIDLTPDYTMESHQRDHENYVSC SQSRRRRAKALLDFERHDDDELGFRKNDII TIISQKDEHCWVGELNGLRGWFPAKFVDIL DERSKEYSVAGDDSVTEGITDLIRGTLSPS IKSIFEHGLKKPSLLGGPCHPWLFIEEAAS REVERDFDSVYSRLVLCKTYRLDEDGKVLT PEELLYRGVQSVNVSHDAAHAQMDVKLRSL ISIGLNEQVLHLWLEVLCSSLPTVEKWYQP WSFLRSPGWVQIKCELRVLSKFAFSLSPDW ELPVKREDKEKKPLKEGVQDMLVKHHLFSW DIDG | *X. laevis* RabGAP-5 (SGSM3, RUTBC3) was expressed as TwinStrep-(linker)-3C-RabGAP-5 fusion and digested with HRV 3C protease. |
| ΔRabex5 | GGGGGSLETDRVSKDFIEFLKTYQKAGHDV YKLSKIFLEAMHHKRESNIDEQSEFTQDFY | Rabex5 truncation mutant without membrane-targeting |

| | | |
|---|---|---|
| | QNTADKLQMYWKVSPDKVEKVMDQIERFIM TRLYKHVFCPETTDDEKKDLTVQKRIRALH WVTLQMLCVPVNEDIAEVSDMVVKAITDII EMDSKRIPRDKLACITRCSKHIFNAIKITK NEPASADDFLPTLIYIVLKANPPRLQSNIQ YITRFCNPSRLMTGEDGYYFTNLCCAVAFI EKLDGQSLNLSEEEFSRYMSGQASPKKQDL ENWPEDTCTGVKQMHRNLDLLTQLSKRQEH IVNGAKKLEKDLIDWTDEVTKEVKDIVEKY PLNIKTASQALAALESENVEDDNLPPPLQP QVYAG | domains (1-131) was expressed as (linker)-TEV-$Gly_4$-ΔRabex5 fusion and digested with TEV(S219V) protease when co-expressed with Rabaptin5 and as $His_6$-(linker)-TEV-$Gly_4$-ΔRabex5 when expressed alone. Also stable in monomeric form. |
| $\Delta_{RBD}$Rabaptin5 | GAAEPGSVVQPDATLQQRVQELERDNAEFL RTKQLLEQEFNQKRAKFKELYLSKEEDLKH QQAVIQVAQEEIVQLNIKLSQAQAEMENIK AVATVSENTKQEAIDEVKKQWQEEVASHQA IMKETVREYELQFHHRLEQERAQWGQYRES VEREIAELRRRLSEGQQEENLENDMKKAQE DAEKLRSVVMPMEKEIGTLKEKLTEAEEKI KDLEASKNQVEEQRTRKRKEKETVQKEEEC RKVILEESLPNLKQEELLNSSHSSIHSLDT DIMLHEGDSFNKQEDLFKDGLRRAQSSDSL GASGPLQTKTLGYNNKAKSAGNLDESDFGP LVGADSVSENFDTSSLGSLHMPSGFMLTKD QEKAIKAMTPEQEETASLLSSVTQTLECTY VPPSDYRLVSETEWNLLQKEVQTAGNKLGR RCDMCSNYEKQLQVIQTQEAEIRDQVKKLQ TMLRQVNDQLEKTLKDKKDLEDYMKQNTEE TSTQISTLTLRINQSETLLADLQQAFHTGK RNIQDQMAVLMHSREQVSEELLRLQRDNES LQGKHSLHVSLQQSEIFNLPETSEELQNLV LKYREDIISVRTATDHLEEKLKAEILFLKE QIQAEQCQKENIEETLQIEIENCKEEMAST SSLQLELDRIKAEREQLEVSLQEKTQELQN LQTLKDSLENQLKKETGSKASLEQLAFEEK NKAQRLQTELDVS | Rabaptin5 mutant with truncated Rab5-binding domains (217-319 and 807-854) was expressed as TwinStrep-(linker)-TEV-Rabaptin5 fusion and digested with TEV(S219V) protease. |

**Table S4. Key reagents and resources used in this study**

| Name | Source | Identifier |
|---|---|---|
| **Bacterial strains** | | |
| BL21(DE3) | New England BioLabs | C2527 |
| BL21(DE3)-RIL [pRK793] | Addgene | 8827 |
| DH10EMBacY | Frederic Garzoni (EMBL Grenoble) | / |
| DH5α | Thermo Fisher Scientific | 18265017 |
| **Insect cell strains** | | |
| HighFive | Thermo Fisher Scientific | B85502 |
| Sf9 | Oxford Expression Technologies | 600100 |
| **Reagents** | | |
| 2-mercaptoethanol | Sigma Aldrich | M6250-100ML |
| Biorad Protein Assay Dye | Biorad | 5000006 |
| catalase | Sigma Aldrich | C40 |
| CF488A maleimide | Sigma Aldrich | SCJ4600016-1UMOL |
| CHAPS hydrate | Sigma Aldrich | C3023-5G |
| CLPETGG peptide | Biomatik | / |
| cOmplete Protease Inhibitor Cocktail | Roche | 5056489001 |
| D-desthiobiotin | Sigma Aldrich | D1411-500MG |
| DiO | Sigma Aldrich | D4292 |
| DMPE-PEG2000 | Avanti Polar lipids | 880150C |
| DNase I | Thermo Fisher Scientific | 90083 |
| DOGS-NTA | Avanti Polar lipids | 790404C |
| DOPC | Avanti Polar lipids | 850375C |
| DOPS | Avanti Polar lipids | 840035C |
| DTT | Sigma Aldrich | D5545, D0632 |
| ESF 921 serum free insect media | Expression Systems | 500304 |
| Express Five SFM (1X) | Thermo Fisher Scientific | 10486-025 |
| GDP | Sigma Aldrich | G7127 |
| glucose oxidase | SERVA | 22778.01 |
| glycerol | Sigma Aldrich | G5516-1L |
| GMP-PNP | Jena Bioscience | NU-401-10 |
| GTP | Jena Bioscience | NU-1012-1G |
| HisPur Ni-NTA resin | Thermo Fisher Scientific | 88221 |
| hydrogen peroxide ($H_2O_2$) | Sigma Aldrich | 216763-500ML-M |
| IGEPAL CA-630 | Sigma Aldrich | I8896-50ML |
| imidazole | Sigma Aldrich | I2399 |

| | | |
|---|---|---|
| Insect GeneJuice | Merck Millipore | 71259-3 |
| IPTG | Bartelt | 6.259 683 |
| L-Glutamine | Thermo Fisher Scientific | 25030024 |
| mant-GDP | Jena Bioscience | NU-204S |
| methyl-β-cyclodextrin | Sigma Aldrich | 332615 |
| Phusion HF DNA polymerase | New England BioLabs | M0530S |
| Pierce BCA Protein Assay Kit | Thermo Fisher Scientific | 23227 |
| PMSF | Sigma Aldrich | P7626 |
| potassium acetate (KOAc) | Sigma Aldrich | P1190-500G |
| Protino Ni-IDA Resin | Macherey Nagel | 745210.3 |
| Q5 HF DNA polymerase | New England BioLabs | M0491S |
| sodium chloride (NaCl) | Sigma Aldrich | S3014 |
| sulfo-Cy5 (sCy5)-maleimide | Lumiprobe | 23380 |
| sulfuric acid ($H_2SO_4$) | Merck Millipore | 1007311000 |
| Taq DNA ligase | New England BioLabs | M0208L |
| TCEP | Sigma Aldrich | C4706 |
| Triton X-100 | Sigma Aldrich | X100-100ML |
| Trolox | Sigma Aldrich | 238813 |
| **Consumables** | | |
| Amicon Ultra-2 concentrators | Millipore | Z740164-24EA |
| black 384-well NBS microplate | Corning | 3820 |
| coverslips 24×50 mm, no. 1.5H | Marienfeld | 107222 |
| LiposoFast 100 nm membrane | Sigma Aldrich | Z373419-50EA |
| Norland optical adhesive 63 | APM Technica | 284239 |
| PD10 desalting columns | GE Healthcare | GE17-0851-011 |
| Pur-a-dialyzer dialysis kit | Sigma Aldrich | purg60010-1kt |
| Vivaspin 20 concentrators | Sartorious | Z614599 |
| Zeba desalt spin columns | Thermo Fisher Scientific | PIER89882 |
| PCR tubes (reaction chambers) | Biozym Scientific | 710920 |
| **Purification columns** | | |
| HiLoad 16/600 Superdex 200 PG | GE Healthcare | 28989335 |
| HiLoad 16/600 Superdex 75 PG | GE Healthcare | 28989333 |
| MonoQ 5/50 GL | GE Healthcare | 17516601 |
| StrepTrap HP 1 mL | GE Healthcare | 28907546 |
| StrepTrap HP 5 mL | GE Healthcare | 28907547 |
| Superdex 200 Increase 10/300 | GE Healthcare | 28990944 |
| Superdex 75 Increase 10/300 | GE Healthcare | 29148721 |
| **Plasmids** | | |
| pet30b-7M SrtA | Addgene | 51141 |

**Mov. S1. CF488A-Rab5 collectively binds the SLB after Rabex5:Rabaptin5 addition (related to Fig. 1B).**

Fluorescently labeled CF488A-Rab5:GDI complex was incubated with free GDI and GTP on glass-supported lipid bilayer in 30 µL reaction buffer. Nucleotide exchange was triggered 10 min after the fluorescence signal equilibrated by 20 µl Rabex5:Rabaptin5 injection (time 0). The fluorescence signal starts accumulating cca. 10 min after GEF addition and correlates with prenylated CF488A-Rab5 SLB binding after activation by the nucleotide exchange. The measured signal saturates after 40 min, indicating the reaction reached steady state. The reaction composition at the final reaction volume was 500 nM CF488A-Rab5:GDI, 2 µM GDI, 50 µM GDP, 500 µM GTP and 200 nM Rabex5:Rabaptin5. The TIRF sequence was acquired on Plan-APOCHROMAT 63x/NA 1.46 immersion objective and 2.5x optovar with 30 s interval. Field of view: 18,2 x 18,2 µm (10 fps).

**Mov. S2. Prenylated CF488A-Rab5 ensemble undergoes collective switching with increasing delay times, depending on the GEF amount (related to Fig. 1C).**

Shown are two CF488A-Rab5 activation reactions at different Rabex5:Rabaptin5 concentrations. Labeled CF488A-Rab5:GDI, free GDI and GTP were incubated on SLB in 30 µL reaction buffer. Nucleotide exchange was triggered 10 min after the fluorescence signal equilibrated by injecting 20 µL of 400 nM (left) or 40 nM (right) Rabex5:Rabaptin5 (time 0). The increase in CF488A-Rab5 fluorescence appeared at two distinct delay times, which is in agreement with stochastic bistable model of Rab5 collective activation. The reaction composition at the final reaction volume was 500 nM CF488A-Rab5:GDI, 2 µM GDI, 50 µM GDP, 500 µM GTP and 400 nM (left) or 40 nM (right) Rabex5:Rabaptin5, respectively. The sequences were acquired on Plan-APOCHROMAT 63x/NA 1.46 immersion objective and 2.5x optovar with 30 s interval. Field of view for respective sequence: 18.4 x 18.4 µm (15 fps).

**Mov. S3. Rabex5:Rabaptin5 is recruited to the SLB during Rab5 activation (related to Fig. 2D).**

Shown are membrane recruitment profiles for CF488A-Rab5 (cyan, left) and Rabex5:sCy5-Rabaptin5 (yellow, right) during Rab5 activation reaction. We added 200 nM of labeled Rabex5:sCy5-Rabaptin5 10 min after CF488A-Rab5:GDI and free GDI mixture equilibrated (time 0). An increase in fluorescence intensity is observed in both channels. The reaction composition at the final reaction volume was 500 nM CF488A-Rab5:GDI, 2 µM GDI, 50 µM GDP, 500 µM GTP and 200 nM Rabex5:sCy5-Rabaptin5. The sequences were acquired on Plan-APOCHROMAT 63x/NA 1.46 immersion objective and 2.5x optovar with 30 s interval. Field of view for respective sequence: 33.2 x 33.2 µm (20 fps).

**Mov. S4. Single particles of sCy5-Rab5 display prolonged lifetimes on SLB surface after nucleotide exchange (related to Fig. 3B).**

Single particle sequences for cCy5-Rab5 before (GDP, left) and after nucleotide exchange with

GEF (GTP, right). **(Left)** Reaction mixture composed of CF488A-Rab5:GDI, GDI and GTP was doped with diluted sCy5-Rab5:GDI and left to equilibrate. Membrane binding of sCy5-Rab5 in equilibrium was then captured with high-speed TIRF acquisition at high laser power in presence of oxygen scavenging system to limit photobleaching. Short sCy5-Rab5 trajectories are visible, representing the free membrane-bound Rab5 population, which is continuously re-extracted by GDI. The reaction composition was 500 nM CF488A-Rab5:GDI, 2 µM GDI, 50 µM GDP, 500 µM GTP, cca. 1 nM sCy5:GDI, 60 mM D-glucose, 0.1 mg/ml glucose oxidase, 0.32 mg/ml catalase, 2 mM Trolox. **(Right)** Reaction mixture composed of CF488A-Rab5:GDI, GDI and GTP was doped with diluted sCy5-Rab5:GDI and left to equilibrate. Then, we triggered the nucleotide exchange by injecting 200 nM Rabex5:Rabaptin5 into the reaction chamber. The single particle trajectories were captured after the activation reaction reached steady state. We included an oxygen scavenging system to limit signal loss due to photobleaching. The final reaction composition was 500 nM CF488A-Rab5:GDI, 2 µM GDI, 50 µM GDP, 500 µM GTP, cca. 50 fM sCy5:GDI, 60 mM D-glucose, 0.1 mg/ml glucose oxidase, 0.32 mg/ml catalase, 2 mM Trolox. The TrackMate-generated particle trajectories are depicted at 20 frame depth in yellow. The sequences were acquired on Plan-APOCHROMAT 63x/NA 1.46 immersion objective and 2.5x optovar with 100 ms interval. Field of view for respective sequence: 52.2 x 52.2 µm (20 fps).

**Mov. S5. Collective CF488A-Rab5 activation progresses as a traveling wave in presence of RabGAP-5 (related to Fig. 4D).**
Fluorescently labeled CF488A-Rab5:GDI complex was incubated with free GDI, GTP and intermediate amounts of RabGAP5 on glass-supported lipid bilayer. We triggered nucleotide exchange by injecting Rabex5:Rabaptin5 and monitored the reaction on SLB plane. We found the collective activation reaction to progress along the SLB as a wave front roughly 90 min after GEF addition. The time denotes relative duration of sequence at selected region of interest, not time after GEF addition. The reaction composition at the final reaction volume was 500 nM CF488A-Rab5:GDI, 2 µM GDI, 50 µM GDP, 500 µM GTP, 50 nM RabGAP-5 and 80 nM Rabex5:Rabaptin5. The TIRF sequence was acquired on Plan-APOCHROMAT 63x/NA 1.46 immersion objective and 1x optovar with 30 s interval. Field of view: 130.6 x 130.6 µm (15 fps).

**Mov. S6. Rab5 activation wave covers large field of view.**
A walk around the activation wave front. CF488A-Rab5:GDI was incubated in presence of free GDI and RabGAP-5 and nucleotide exchange was triggered by Rabex5:Rabaptin5 addition. **(Left)** A distinct activation wave formed after more than an hour. The time denotes relative duration of the walk sequence across the region of interest, not time after GEF addition. **(Right)** Track of the field of view movements in the movie sequence. The respective field of view is shown in red. The reaction composition at the final reaction volume was 500 nM CF488A-Rab5:GDI, 2 µM GDI, 50 µM GDP, 500 µM GTP, 50 nM RabGAP-5 and 80 nM Rabex5:Rabaptin5. The TIRF sequence was acquired on Plan-APOCHROMAT 63x/NA 1.46 immersion objective and 1x optovar with 0.5 s interval, 15 ms exposure time and 100 EM gain. Field of view: 89.9 x 89.9 µm (20 fps).
